# Variation in morpho-physiological and metabolic responses to low nitrogen stress across the sorghum association panel

**DOI:** 10.1101/2022.06.08.495271

**Authors:** Marcin W. Grzybowski, Mackenzie Zwiener, Hongyu Jin, Nuwan K. Wijewardane, Abbas Atefi, Michael J. Naldrett, Sophie Alvarez, Yufeng Ge, James C. Schnable

**Affiliations:** Center for Plant Science Innovation, University of Nebraska-Lincoln, Lincoln, NE USA; Department of Agronomy and Horticulture, University of Nebraska-Lincoln, Lincoln, NE USA; Department of Plant Molecular Ecophysiology, Institute of Plant Experimental Biology and Biotechnology, Faculty of Biology, University of Warsaw, Warsaw, Poland; Department of Biological Systems Engineering, University of Nebraska-Lincoln, Lincoln, NE USA; Department of Agricultural and Biological Engineering, Mississippi State University, Starkville, MS USA; California Strawberry Commission, San Luis Obispo, CA USA; Proteomics and Metabolomics Facility, Nebraska Center for Biotechnology, University of Nebraska-Lincoln, Lincoln, NE USA

## Abstract

Access to biologically available nitrogen is a key constraint on plant growth in both natural and agricultural settings. Variation in tolerance to nitrogen deficit stress and productivity in nitrogen limited conditions exists both within and between plant species. Here we quantified variation in the metabolic, physiological, and morphological responses of a sorghum association panel assembled to represent global genetic diversity to long term, moderate, nitrogen deficit stress and the relationship of these responses to grain yield under both conditions. Grain yield exhibits substantial genotype by environment interaction while many other morphological and physiological traits exhibited consistent responses to nitrogen stress across the population. Large scale nontargeted metabolic profiling for a subset of lines in both conditions identified a range of metabolic responses to long term nitrogen deficit stress as well as several metabolites associated with variation in the degree of yield plasticity specific sorghum genotypes exhibited in response to nitrogen deficit stress.

## Introduction

Malthus predicted that exponential population growth would always surpass linear increases in food production resulting on constant famine (Malthus, 1798). Both dramatic increases in total agricultural land and technological innovations have staved off Malthusian catastrophy in the 20th and early 21st century. One of the key technological innovations was invention and widespread adoption of the Haber–Bosch process, which reduces atmospheric nitrogen gas (N_2_) to reactive forms of N, to provide an abundance and reliable source of nitrogen fertilizer for agriculture (Erisman *et al*., 2008). The widespread adoption of synthetic nitrogen fertilizers have dramatically increased crop yields but these increases have not come without some negative externalities, including increased greenhouse gas emissions and decreases in rural water quality (Zhang *et al*., 2015). In addition, for many non-irrigated agricultural systems the cost of fertilizer is the either single biggest variable input cost of production, or the second biggest after the cost of seed (Rothstein, 2007). As the human population continues to grow and populations around the world shift to more calorie intensive diets, incentives and pressure to agricultural productivity will increase as well (Foley *et al*., 2011; Ramankutty *et al*., 2018). It has been estimated that only 30-40% of nitrogen fertilizer is taken up and utilized by crops (Raun and Johnson, 1999). Increasing the nitrogen use efficiency of major agricultural crops would enable farmers to meet these growing requirements for food production with stable or decreasing applications of nitrogen fertilizer, increasing farmer profitability will decrease the environmental and energy footprint of agriculture (Hakeem *et al*., 2011).

Substantial genetic variation in nitrogen use efficiency exists within crop plants (Cañas *et al*., 2012; Liu *et al*., 2021). Between 1969 and 2010 European wheat breeders increased the nitrogen use efficiency of wheat by an estimated one third of one percent per year (Cormier *et al*., 2013). The global impact of a 1% increase in nitrogen use efficiency is estimated to be $1 billion dollars per year (Kant *et al*., 2011). Understanding the genes controlling variation in nitrogen use efficiency and the other phenotypes associated with these differences would aid in both evaluating the feasibility of increasing nitrogen use efficiency in different crops – while sustaining the high yields necessary to meet global demand for food – and, where feasible, designing breeding strategies to achieve such an increase. However, nitrogen use efficiency is a complex trait and multiple morpho-physiological and metabolic mechanisms likely play roles in determining how well or poorly a given plant genotype can compensate for limited N availability in different environments and at different life stages. Understanding the morpho-physiological and metabolic mechanisms associated with differences in tolerance for nitrogen deficit stress in agriculturally relevant environments represents a stepping stone to the subsequent identification of genetic loci and finally to crop improvement via breeding or engineering. To date the majority of research on the morpho-physiological and metabolic responses of plants to nitrogen deficit stress has been conducted in controlled environment conditions, particularly emphasizing severe stress applied early in development (Amiour *et al*., 2012; Banerjee *et al*., 2020; Gao *et al*., 2015). The nitrogen deficit stress experienced by crops in agricultural settings is typically less extreme and may not produce obvious visual effects, but is sufficient to result in substantial grain or biomass decrease over the course of a growing season. Collecting phenotypic and metabolic data from large sets of genotypes experiencing agriculturally relevant degrees of stress under field conditions can provide substantial insight into natural variations in stress response and tolerance within individual crop species (Obata *et al*., 2015).

Here we quantified crop yield and eight morpho-physiological traits from a large and diverse sorghum population (*Sorghum bicolor* L.) grown to maturity in field conditions under both nitrogen limiting and non nitrogen limiting conditions. For a subset of 24 replicated genotypes, large scale metabolic profiling was conducted from leaf tissue collected at the flowering stage. Significant plasticity and genotype x environment interactions were observed for both yield and a subset of metabolic traits, while substantially less genotype x environment interaction was observed for morpho-physiological traits. The abundance of several metabolites at flowering exhibited significant correlations with plant performance (e.g. yield) at maturity.

## Material and methods

### Field experiment, and phenotypic data and tissue collection

A replicated field trial was planted at the University of Nebraska-Lincoln’s Havelock Farm Location (N 40.861, W 96.598) on June 08, 2020. The experiment was laid out in a RBCD design, initially with three blocks each under sufficient nitrogen (80 lbs/acre) and low nitrogen (no supplemental nitrogen) treatment conditions and 416 plots per block, including 347 genotypes from the sorghum association panel (Casa *et al*., 2008), and BTx623 as a repeated check. Each plot consisted of a single 2.3 meter row of plants from a single genotype, with 0.76 meter spacing between parallel and sequential rows.

A mixture of hand measured traits and traits predicted from hyperspectral data (see below) were employed to assess the response of sorghum to nitrogen deficit stress. The date of flowering for each plot was scored when 50% of surviving plants had reached anthesis. Plant height was measured from the soil surface to the flag leaf collar after flowering. Panicles per plot were hand-counted. One to three panicles per plot were hand harvested from each plot, dried, and threshed and the resulting grain weighed. Grain weight per panicle was multiplied by panicles per plot to estimate yield per plot.

Between 5 and 12 August 2020, hyperspectral reflectance data was collected from the second leaf from top of the plant from single plant per block using a FieldSpec4 (Malvern Panalytical Ltd., formerly Analytical Spectral Devices), following the protocol outlined in Ge *et al*. (2019). A set of 265 leaf samples (130 from HN and 135 from LN) were selected for ground truth measurements. Leaf chlorophyll concentration (CHL) was measured with a handheld chlorophyll concentration meter (MC-100, Apogee Instruments, Inc., Logan, UT), and leaf area (LA) was measured with a leaf area meter (LI-3100, LI-COR Biosciences, Lincoln, NE). Next, samples were placed in a oven set to 50°C and dried over 72 h. Dry weight (DW) of the leaves was then recorded with digital balance. Specific Leaf Area (SLA, m^2^/kg) was calculated as LA/DW. Finally, dried plant leaves were sent to commercial lab (Ward Laboratories, Inc., Kearney, NE) where the samples were ground, homogenized, and analyzed for analysis of nutrient content: nitrogen, potassium and phosphorus.

For 96 plots representing 24 genotypes replicated in two blocks each under sufficient and low nitrogen treatments, leaf tissue was collected for metabolomics analysis. For each plot, a single plant was selected, avoiding edge plants where possible. From this plant eight leaf punches of 0.33 cm^2^ in area were collected from the middle section of the leaf below the flag leaf (e.g the penultimate leaf) and immediately frozen in liquid nitrogen. Samples were collected between 9:00 AM and 1:00 PM on August 12 2020.

### Modeling traits based on hyperspectral data

Five models were developed to predict chlorophyll, nitrogen, phosphorus, and potassium concentration as well as specific leaf area from hyperspectral reflectance data, following the approach described in Ge *et al*. (2019). Measured intensity values for each wavelength were zero centered and scaled to unit variance. Wavelengths below 450 nm and above 2400 nm were discarded. Predictive models were built separately for each trait using partial least squares regression implemented in the *pls* v.2.8.0 (Liland *et al*., 2021) and *caret* (Kuhn, 2008). Prior to the modeling, data were split into training (n=185) and validation set (n=80). This was done to avoid the risk of misleadingly high prediction accuracy resulting from over fitting. Decisions regarding model tuning and performance evaluation were made based on root mean squared error (RMSE) of five-fold cross validation using training set (n=185). After final models were trained, their performance was evaluated using the validation set (n=80). Final models were applied to equivalently zero centered and scaled hyperspectral reflectance measurements collected from the remaining sorghum plots.

### Untargeted metabolomics using LC-MS/MS

Samples were extracted using cold methanol:acetonitrile (50:50, v/v) spiked with 100 M of CUDA (12-[(cyclohexylcarbamoyl)amino]dodecanoic acid). The tissue samples were disrupted and homogenized by adding 2 stainless steel beads (SSB 32) using the TissueLyserII (Qiagen) at 20 Hz for 5 mins. After centrifugation at 16,000 g, the supernatants were collected and the same extraction was repeated on the pellet one more time. The supernatants were pooled and vacuum dried down using a SAVANT speed-vac. The pellets were re-dissolved in 100 *µ*L of 30% methanol. Blank tubes were extracted alongside the samples to remove contaminant background from the data analysis. In addition, an aliquot of the samples was pooled to make a quality control (QC) sample which was run between every 10 samples in order to correct for batch effect. Two separate LC-MS/MS workflows running on a Thermo Vanquish LC system interfaced with a Thermo QE-HF mass spectrometer were used to profile the metabolites. For the hydrophobic compounds, a ACCQ-TAG ULTRA C18 column (1.7 *µ*m, 2.1 mm × 100 mm, Waters) was used flowing at 0.3 mL/min at 40 °C. The gradient of the mobile phases A (0.1% formic acid in water) and B (0.1% formic acid in acetonitrile) was as follow: 2% B for 2 min, to 50% B in 11 min, to 90% B in 2 min, hold at 90% B for 1 min, to 2% B in 0.5 min. The QE-HF was run in a data-dependent acquisition mode triggering on single charge peaks using a mass range of 67 to 1000 m/z at 60,000 resolution, with an AGC target of 3e6 and a maximum ion time of 100 ms for both positive and negative ion scans. The isolated ions were further fragmented by HCD using isolation window of 1.6 m/z and scanned at a resolution of 15,000. For the polar compounds, a XBridge Amide 3.5(4.6 × 100 mm, Waters) was used flowing at 0.4 mL/min at 45 °C. The gradient of the mobile phases A (10 mM ammonium formate/0.125% formic acid in water) and B (10 mM ammonium formate/0.125 formic acid in 95% acetonitrile) was as follow: 100% B for 2 min, to 70% B in 5.7 min, to 40% B in 1.8 min, to 30% in 0.75 min, to 100% B in 2.5 min. The QE-HF was run in a data-dependent acquisition mode triggering on single charge peaks using a mass range of 60 to 900 m/z at 60,000 resolution, with an AGC target of 1e6 and a maximum ion time of 100 ms for both positive and negative ion scans. The isolated ions were further fragmented by HCD using isolation window of 1.6 m/z and scanned at a resolution of 15,000.

### LC-MS/MS data analysis

Data from LC-MS/MS analysis were process with MS-Dial software v4.70 for peak detection, deconvolution, alignment, quantification, normalization, and identification (Tsugawa *et al*., 2015). Background peaks detected in blank extracts were filtered out. Intensity drift was corrected using the local regression (LOESS) for QC batch normalization, and zero intensities were replaced by 10% of the minimum peak height. The identification was done using the curated mass spectral public libraries (http://prime.psc.riken.jp/compms/msdial) for MS/MS positive (290,915 entries, April 2021) and MS/MS negative (36,848 entries, April 2021). Metabolites missing in more than 80% of the total samples were removed. The remaining 3,496 metabolites from all four analytical conditions were manually checked for Gaussian chromatographic peak and, peak alignment and MS/MS profile. Identified metabolites were classified either as level I when peak matched to m/z and retention from an in-house library prepared from authentic standards, or as level II based on their spectral similarities with public/commercial spectral libraries in accordance with the Metabolomics Standards Initiative guidelines (Sumner *et al*., 2007).

### Phenotypic data analysis

All statistical analyses were conducted in R v.4.1.2 (R Core Team, 2021). The meta-package *tidyverse* v.1.3.1 was employed for data processing and visualization (Wickham *et al*., 2019). In order to analyze the impact of the treatment effect on morpho-physiological traits and metabolites, mix-models were fit to each trait – after being transformed using the Box-Cox method – using the *lmer* function provided by the *lme4* package (Bates *et al*., 2015). The full model fit contained the treatment as fix effect and genotype as random effect, wheres reduced model only genotype. The difference between this two models were evaluated using the likelihood ratio test (LRT) to obtain p-values for the significance of treatment effects. P-values from metabolite data analysis were corrected for multiple tests using false discovery rate (FDR) (Benjamini and Hochberg, 1995), and values below 0.05 were considered to be statistically significant.

A more complex model which, in addition to treatment (nitrogen) as a fixed effect and genotype as a random effect, also included genotype by environment (GxE) interaction as random effects was fit for each metabolite in order to estimate total variance potentially explainable by each of these three factors (Brommer, 2013; de Jong *et al*., 2019). Metabolites for which variance estimated from the model for one or more parameters were zero, or close to zero, and therefore singular fit of the model was obtain, were excluded for analysis.

Broad-sense heritabilities were estimated from the following equation:

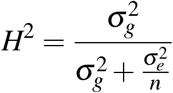

where *σ* ^2^_g_ is genetic variance, *σ* ^2^_e_ is residual variance, and n is the number of replicates. Variances were obtain from mixed model fitted separately to values from each experimental conditions with genotypes treated as random effect.

Principle component analysis for metabolite values across the 96 samples were calculated using the *PCA* function provided by the *FactoMineR* package (Lê *et al*., 2008). Pearson correlation analysis between yield and metabolites were done with *cor*.*test* function in R.

Yield predictions were done based on three metabolite data sets: all identified metabolites (n=3,496), metabolites with confident annotation (n=145), and the same number of metabolites with unknown annotation (n=145). Analysis were done with caret framework (Kuhn, 2008). Random forest were fitted with *ranger* package (Wright and Ziegler, 2017) and elastic-net regression with *glmnet* package (Friedman *et al*., 2010). Prior to the analysis, yield and metabolites values were Box-Cox transformed and scaled with *preProcess* function. Repeated 100x times five-fold cross validation were used to determinate optimal parameters for each model based on minimization root mean square error (RMSE). Importance value were calculated with *varImp* function from *caret* package based on permutation. The mean squared error is computed on the out-of-bag data for each model, and then the same computed after permuting a single variable. The differences are averaged and normalized by the standard error and scaled to values between 0 and 100.

## Results

### Genetic variability of sorghum’s response to differential nitrogen application

A population of 347 sorghum genotypes drawn from the Sorghum Association Panel (SAP) (Casa *et al*., 2008) were grown under two nitrogen treatments with replication in Lincoln, Nebraska: low nitrogen (LN; no supplemental nitrogen) and high nitrogen (HN; 90 kg/ha, following local agronomic recommendations to avoid nitrogen limitations on yield in sorghum). A mixture of manually scored – leaf number, flag leaf length, flag leaf width, plant height, days to flowering– and phenotypes estimated from hyperspectral reflectance data – specific leaf area (SLA), chlorophyll content (CHL) and nitrogen (N), phosphorus (P) and potassium (K) content following previous workflow (Ge *et al*., 2019) – were collected from plants grown under both conditions (Table S1).The overall hyperspectral reflectance profile of sorghum leaves collected from plants grown in HN and LN treatments was similar (Fig. S1a), and neither of the first two principle components clearly separated the two treatments (Fig. S1b). Ground truth data were obtained for five traits: CHL, SLA, N, P, and K content from 265 samples, and partial least squares regression (PLSR) were used to predict values for scored six traits for whole panel based on hyperspectral data. Raw spectral data used for PLSR model building, as well as prediction traits were provided in Table S2. Employing five-fold cross validation with ground truth samples the accuracy (R^2^) of phenotypes estimated from hyperspectral reflectance data varied from 0.18 for P to 0.82 for CHL (Table S3). Similar performance was observed in the validation set (n=80, Table S3 and Fig. S2) indicating models were not overfit. Prediction accuracy for K and P were low (R^2^ in validation set 0.22 and 0.25; Table S3), and these traits were excluded from downstream analyses.

The effect of N treatment was statistically significant for all traits evaluated, except for plant height (p < 0.05; likelihood ratio test (LRT); Fig. 1). Flag leaf width and flag leaf length were reduced by approximately 3.5% and 6.5% respectively under LN treatment. Plants grown under LN took 4% more time to flower. Larger differences were observed in chlorophyll and nitrogen content, with reductions of 15.3% and 13.8% respectively under LN treatment. However, the single largest impact of low nitrogen stress was observed on grain product, with a 48% reduction in grain yield under LN treatment.

**Figure 1.**
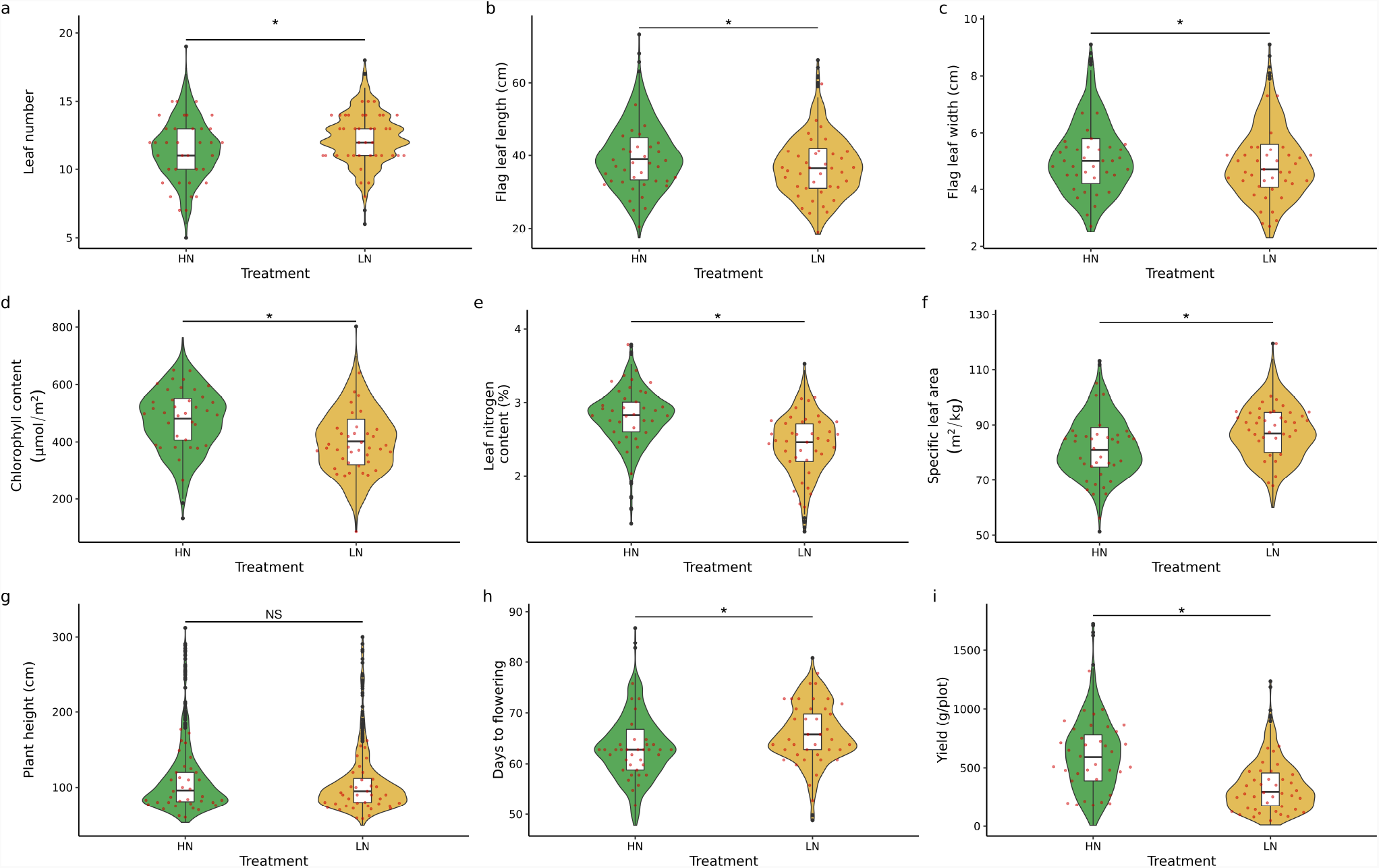
Phenotypic difference of morpho-physiological traits across two treatment conditions. Statistical significance of N treatment were determined by likelihood ratio test (LRT) on mix model with treatment denote as fix effect and genotype as random. Asterisks indicate p-value < 0.05. Red dots indicated values for genotypes selected for metabolomics analysis. HN - high nitrogen, LN - low nitrogen.

While overall population level responses to nitrogen deficit treatment were statistically robust, individual genotypes often exhibited different degrees of response to nitrogen treatment. A mixed model, considering genotype, treatment, and the interaction between genotype and treatment (genotype-by-environment, GxE) effects, was fit to each individual phenotypic dataset. A majority of the total variation in plant height and flowering time was explained by differences between genotypes (~ 91% and ~ 85% respectively together on HN and LN; Fig. 2a). In the case of plant height none of variance was attributed to treatment or genotype by environment interaction. For flowering time, only ~ 4% of variance were explain by treatment effect and ~ 3% by GxE. The high degree of genetic control and low GxE effect is reflected in the high degree genetic correlation across treatment conditions for these traits: 0.86 for plant height and 0.8 for flowering time (Fig. S3). Variance in traits related to leaf (leaf number, leaf width, and leaf length) were also mostly explained by genetic factors (> 60% for each of these three traits). However, proportion of variance not explained by any of the factors in the model (e.g. the residual) was substantially greater for each of the three leaf related traits compared to plant height and flowering time, leading to lower correlations across treatment (~ 0.6; Fig. S3). Traits estimated from hyperspectral data (CHL, N, SLA) were comparatively much more plastic across environments (Fig. 2a), but only modest amounts of variance was attributed to GxE for each of these traits. One explanation for this, is fact, that although our PLSR models were accurate (R^2^ > 0.6), it might be still not sufficient to precisely capture GxE.

**Figure 2.**
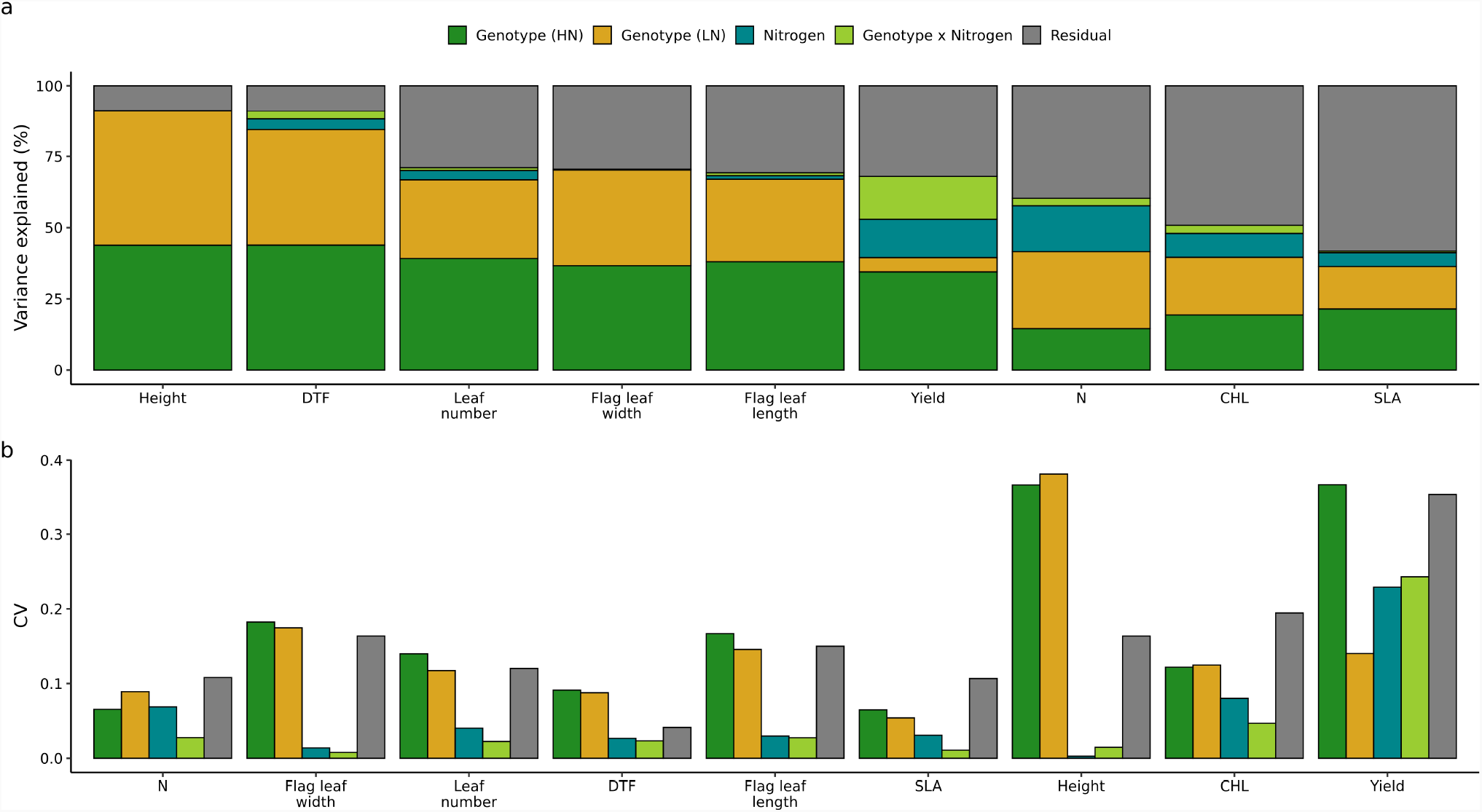
Components of traits variation. **a** shows the proportion of variance attributed to each component for each trait. **b** shows the magnitude of this variance relative to each trait’s mean, using the coefficient of variation (CV; the estimated variance divided by the squared mean of the respective trait). HN - high nitrogen, LN - low nitrogen.

Extensive plasticity of grain yield in response to nitrogen deficit stress was observed across the study population. Among analyzed traits, grain yield exhibited by far the largest proportion of variance attributable to GxE (Fig. 2a), resulting in only moderate genetic correlation between treatment (0.42, Fig. S2). Genotypes with high grain yield in the HN treatment tended to be somewhat more sensitive to low nitrogen stress than genotypes with low grain yield, even under the HN treatment (Fig. S4). However, the correlation between the responses of grain yield to nitrogen deficit stress and grain yield under HN was modest (~ −0.3; Fig. S4). This reflects the relatively large GxE effect of nitrogen treatment on yield. Although highly yielding genotypes on HN are more sensitive to LN stress, this reaction is not consistent and yield of some genotypes are less affected by LN stress.

Coefficients of variation were calculated for each variance component for each trait, following the approach described in de Jong *et al*. (2019). Plant height exhibited the largest relative variation, particularly variation attributed to genetic factors (Fig. 2b), likely reflecting the effects of multiple large effect dwarfing genes segregating for functionally distinct alleles among the lines of the sorghum association panel (Thurber *et al*., 2013). The second largest relative variance was observed for grain yield, in particular for genetic factor under HN. However, relative variance from treatment conditions and GxE were also large, and in fact larger than the variance for any component among the remaining traits.

### Metabolomic changes in sorghum leaves under long-term low nitrogen stress

As morpho-physiological traits scored in this study did not appear to explain the plasticity of sorghum grain yield across different nitrogen availability treatments, we next sought to characterize the responses of a large suite of metabolic phenotypes to differential nitrogen availability in the adult leaves across a subset of sorghum genotypes of the SAP. A set of 24 genotypes were selected to represent the phenotypic and genetic diversity of the SAP (Fig. 1, S5; (Miao *et al*., 2020)). Sampling was timed to coincide with anthesis, with a total of 96 leaf samples collected from two independent plots per genotype per treatment. Each sample was quantified via liquid chromatography - high-resolution mass spectrometry (LC-HRMS) analysis. In order to maximize the number of metabolites detected and quantified each sample was analyzed using both RP (reverse phase) and HILIC (hydrophilic interaction liquid chromatography) separations in both positive and negative ion mode, resulting in the detection and quantification of 115,782 mass spectral features. After filtering out features that were detected in less than 80% of samples, and further manual quality control (as described in the methods section), the number of features was reduced to 3,496, of which 145 could be assigned high confidence annotations (Table S4).

No obvious differences were observed in the distribution of estimated abundance values for high confidence metabolites (n = 145) between HN and LN conditions (Fig. 3a). Samples collected from plants grown in HN or LN were not clearly separated by either of the first two principal components of variation for the abundance of this set of high confidence annotated metabolites although samples collected from plants grown in LN exhibited a tighter distribution of PC1 values than did samples collected from plants grown in HN (Fig. 3b). After correcting for multiple testing via false discovery rate (FDR, (Benjamini and Hochberg, 1995)), the abundance of 62 metabolites changed significantly between samples collected from plants grown in HN or LN (FDR < 0.05). Thirty-four metabolites were more abundant in samples collected from plants grown under HN and 28 more abundant in samples collected from plants grown under LN (Fig. 3c). Although the vast majority of these changes, despite being statistically significant, were relatively modest with less than a two fold change in abundance between treatments. The majority of observed amino acids (17/33) were significantly more abundant in samples collected from plants grown under HN but the amino acids acetylcarnitine and L-carnitine were significantly more abundant in samples collected from plants grown under LN (Fig. 3d). In contrast, half of the phenolic compounds confidently identified in this dataset, such as the eight flavonoids, were significantly more abundant in samples collected from plants grown under LN (Table S4).

**Figure 3.**
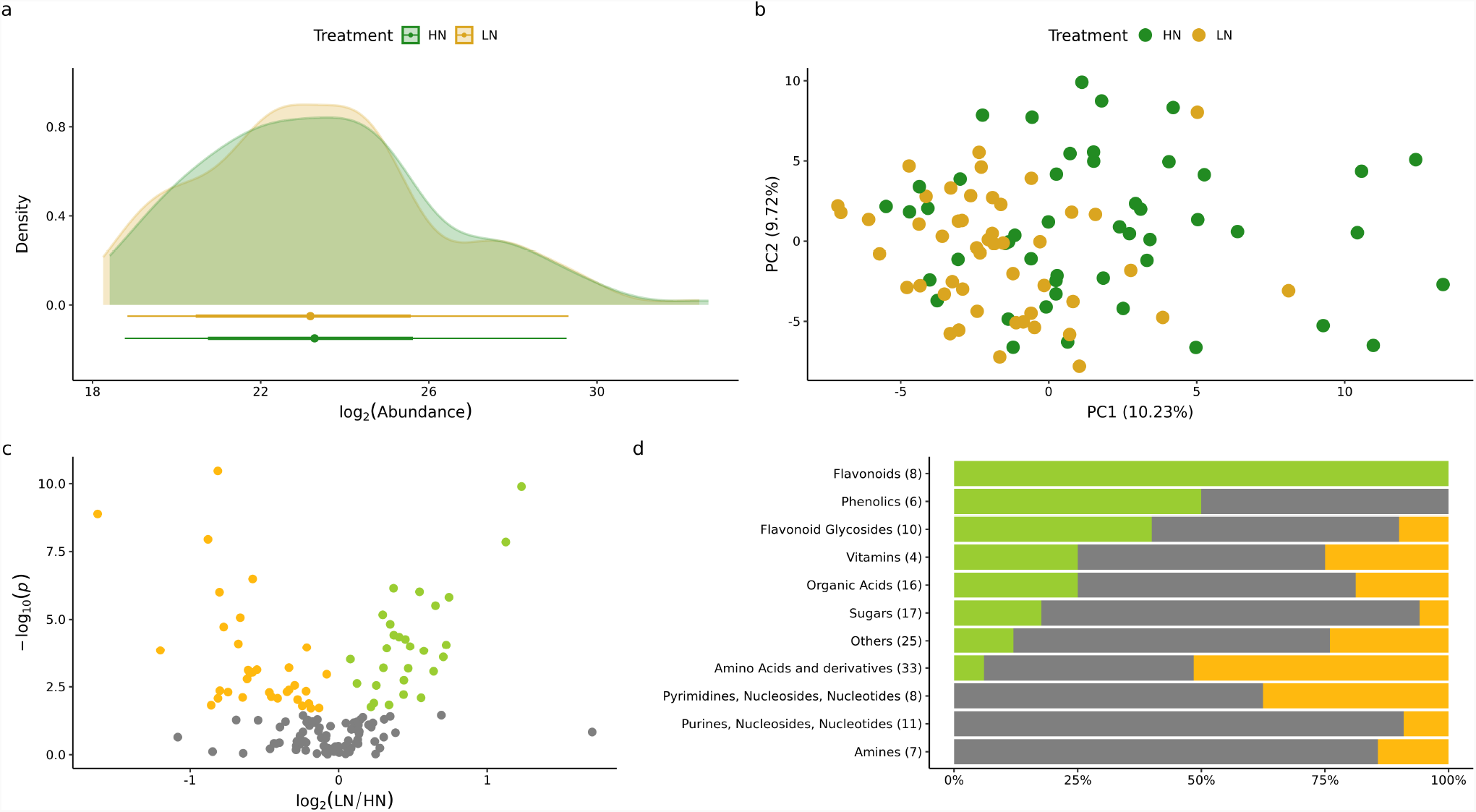
Metabolomics profiling in 24 sorghum genotypes across two nitrogen conditions based on 145 confidently annotated metabolites. **a** Distribution of the 145 confidently annotated metabolites across two treatment conditions. **b** First two principle component (PC) from PCA. Values in bracket indicate amount of variance explained by each component. **c** Volcano plot showing the down regulated (yellow) and up regulated (green) metabolites under low nitrogen (LN) conditions compare to high nitrogen (HN). **d** Proportions of the metabolites with know structures more abundant in samples collected from plants grown under HN (green), more abundant in samples collected from plants grown under LN (yellow), and unchanged (gray).

Similar results to those observed with the set of annotated metabolites were observed when analysed all 3,496 identified mass features. Overall abundance of those compounds was similar across treatment (Fig. S6a-b). Although 337 compounds were significantly different across two treatment condition (FDR < 0.05; Fig. S6c), those changes were rather small, with only 28 compounds being changed larger than two fold between treatments. Finally, PCA based on 3,496 mass features didn’t separate two treatment conditions (Fig. S6d). Interestingly, many of these unidentified metabolites showed relatively high heritability, with mean value 0.6 under HN and 0.68 under LN (Fig. S7). This suggests that natural variation in the contents of these compounds is genetically controlled, which makes a good prospect for furthering their identification and uncovering their biological meaning through genetic studies. In case of known metabolites, variation in the abundance of individual flavonoid and flavonoid glycosides compounds tended to be the most heritable across independent field plots of the same genotype grown in the same environment (Fig. S8).

A similar variance partitioning strategy to that employed for morpho-physiological traits was used to partition variance for each annotated metabolite. For each metabolite a mixed model was fit, including terms for genotypes (genetic effects), differences between N treatments (environmental effects), and genetic differences in the degree of response to N supply (genotype-by-environment, GxE). Likely as a result of the much smaller overall number of datapoints for each metabolic trait relative to each morpho-physiological traits, this model could only be successfully fit for 46 of 145 metabolites. Differences between genotypes typically explained around half of the variance for different metabolites (~ 28% on HN and ~ 31% on LN), while the variance explained by environmental factor was much lower ~ 2% (Fig. 4). The GxE effect explained on average of ~ 7% of variance across the 46 metabolites where a mixed model was successfully fit. Despite the fact, that this value was not very high, it was higher than the average variance explained by the GxE effect for morpho-physiological traits (~1%). The coefficient of variation for each variable for each metabolite vary, but no clear pattern can be observed across different classes of metabolites (Fig. S9). A wide range of different patterns are exhibited by individual metabolites in response to LN stress across different genotypes. Glucose and sucrose both belong to the set of 83 metabolites which did not show any statistically significant differences in abundance between samples collected from plants grown in high nitrogen and plants grown in low nitrogen but which do exhibit consistent patterns of difference in abundance between genotypes across treatments (Fig. 5a-b). Serine, one the amino acids with a statistically significant difference in abundance between samples collected from plants grown in HN and LN exhibits a consistent decreased in abundance across genotypes with ~ 15% of variance explained by the environmental factor (Fig. 5c). Glutamic acid and allantonin both exhibited large GxE effects of ~ 15% and ~ 30% variance for these two metabolites explained by GxE respectively (Fig. 5d-e). Genotypes with comparatively high glutamic acid content in HN saw larger reductions in glutamic acid content in LN. Genotypes with comparatively lower high glutamic acid content in HN saw smaller reductions in LN. Previous study found a decrease in salicylic acid content under low nitrogen stress in sorghum root (Sheflin *et al*., 2019). Here we found increase in salicylic acid content of sorghum leaves under LN (Fig. 5f). This response is consistent across majority of genotypes, although strength of this reaction slightly vary, with genotypes with low salicylic acid content under HN indicating a higher increase under LN.

**Figure 4.**
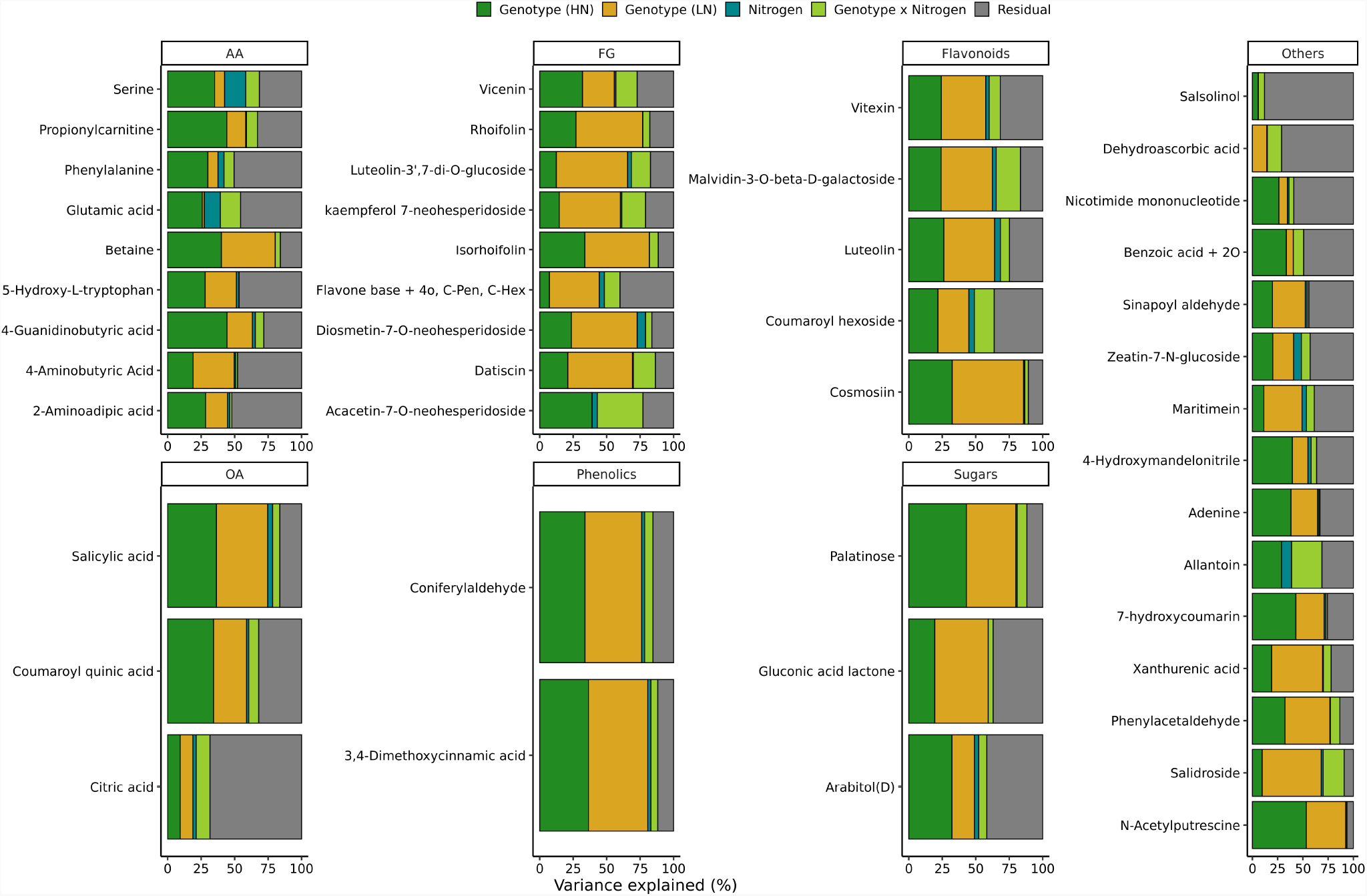
Proportion of variance attributed to each component for each metabolite. HN - high nitrogen, LN - low nitrogen.

**Figure 5.**
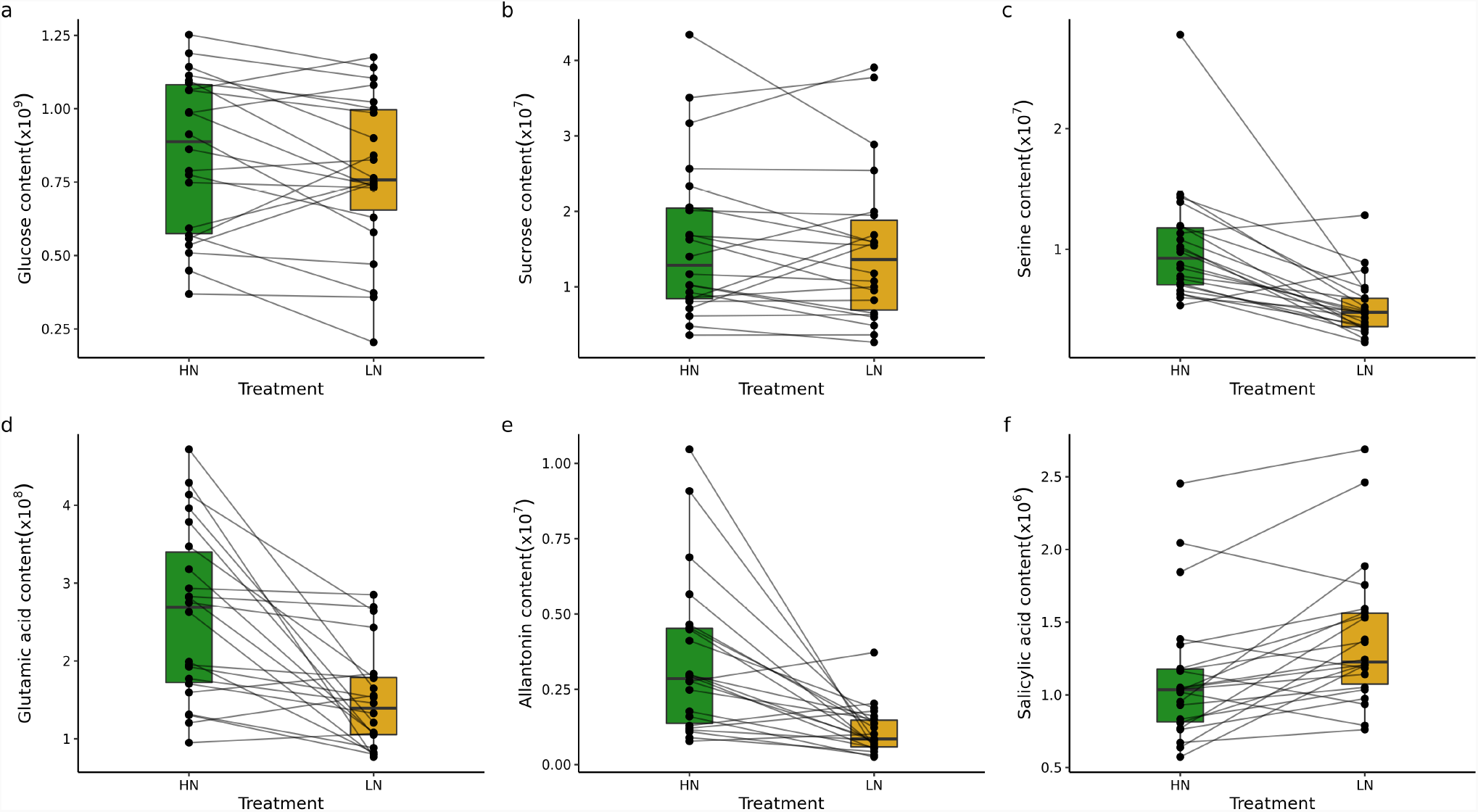
Examples of unchanged **a**-**b**, changed but non-plastic **c** and plastic **d**-**f** metabolites. Each dot indicate genotypic mean and lines connect the same genotype across two treatment conditions. HN - high nitrogen, LN - low nitrogen.

### Correlation between metabolites and yield

The abundance of metabolites was correlated to some degree with observed grain yield values from the same plots (Fig. 6; Table S5). The correlation coefficients between metabolite abundance and grain yield in individual environments (HN or LN) were positively correlated with each other (r = 0.36, p < 0.05; Fig. 6). However, in only a modest number of cases where the correlations between the abundance of individual metabolites and grain yield statistically significant including six metabolites in HN and eleven in LN. Five metabolites were statistically significantly correlated with grain yield in both environments: 4-hydroxymandelonitrile, aconitic acid, ascorbic acid, benzamide and glucose. The strength of the correlations between grain yield and metabolite abundance where relatively modest (r < 0.6) even for those metabolites where statistically significant relationships were observed.

**Figure 6.**
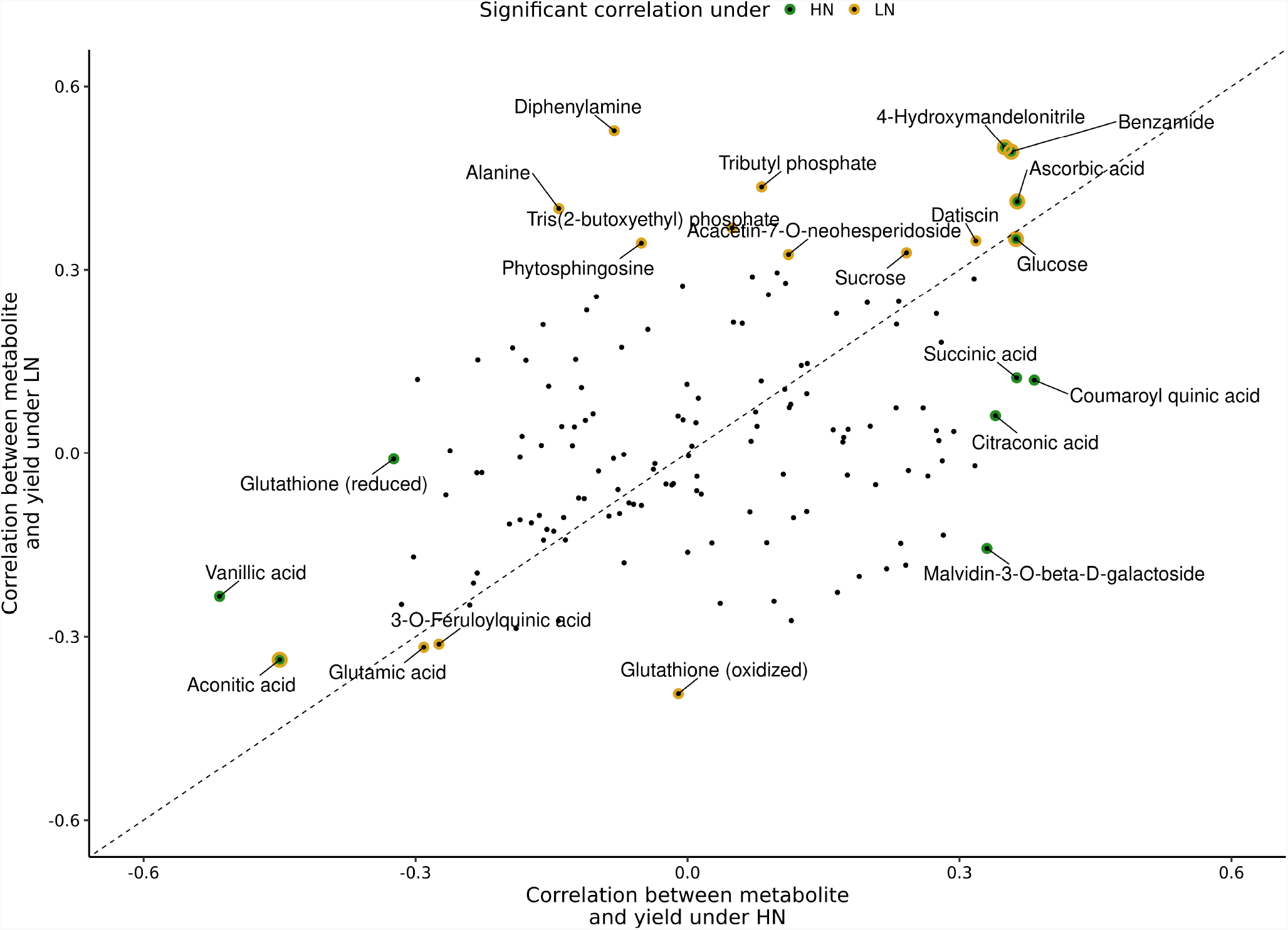
Scatter plot of correlation values of each metabolite and yield in given conditions. Marked metabolites indicate significantly correlated metabolites from Pearson analysis (p < 0.05). HN - high nitrogen, LN - low nitrogen.

Two machine learning approaches, elastic-net regression (GLMNET) and random forest (RF), were evaluated for their potential to predict variation in plot level grain yield from combined metabolite abundance data. Three sets of input data were evaluated with each of the two machine learning approaches. First, the set of all detected metabolites (n=3,496). Second, the set of 145 metabolites with confident annotations. Finally, a set of 145 metabolites selected randomly from the complete set of 3,496 detected metabolites. Both algorithms achieved moderate prediction accuracy however, the accuracy of their predictions was either equivalent to or only modestly exceeded, the prediction accuracy of a simple linear model fit to only the treatment effect, which was able to predict 29% of the total variance in sorghum grain yield data (Fig. S10). Ascorbic acid showed the greatest contribution to the accuracy of the GLMNET model (Fig. S10b) and the third largest contribution with RF (Fig. S10c) in permutation based estimates of feature importance, consistent with the significant correlation between the abundance of this metabolite and grain yield in both conditions (Fig. 6).

## Discussion

Natural variation in tolerance to nitrogen deficient growing conditions has been widely studied in both crops and other plant species. However, the majority of these studies occurred in controlled environments and imposed substantial nutrient deficits that produced visible phenotypic responses even at seedling stages. Here we examined natural variation in both the morpho-physiological and metabolomic impact of long term low intensity nitrogen deficit at a level sufficient to alter grain yield and fitness in sorghum but which does not produce obvious visible stress symptoms.

Grain yield is a complex phenotype that is determined by a number of different component phenotypes (e.g. yield component traits). In arabidopis branching number is correlated with yield and previous study found large plasticity of this trait in response to low nitrogen stress (de Jong *et al*., 2019). In case of rice, various traits such as tiller number, grain number per penile, or 1,000 - grain weight are associated with yield. Interestingly, only tiller number were affected by low nitrogen stress (Liu *et al*., 2021). Finally, in case of maize 1,000 - kernel weight were also not affected by low nitrogen, but substantial decreases in kernel number per cob were observed (Amiour *et al*., 2012). This observation highlights the complexity of how plant yield can be affected by low nitrogen stress. In this study grain weight per panicle was used to represent sorghum yield, and consistently with research done on arabidopsis (de Jong *et al*., 2019), large plasticity in response to low nitrogen stress was observed in this trait.

While grain yield decreased substantially under nitrogen limited conditions for the vast majority of sorghum genotypes, rank order grain yield under high nitrogen conditions was only modestly correlated with rank order grain yield under low nitrogen conditions (Spearman correlation = 0.44; Fig. S3). This suggests efforts to increase grain yield under nitrogen limited conditions will require separate field trials, evaluations and selections from breeding efforts to increase grain yield under non-nitrogen limited conditions. Yield under non-nitrogen limited conditions was negatively correlated with the size of the decrease in yield observed when nitrogen was limited (~ −0.3; Fig. S4). However, because of large GxE effect, this reduction is not consistent across highly yielding genotypes. Some of the reductions are characterized by relatively low loss in yield under low nitrogen conditions. This indicates that it should be possible to produce varieties not only with high yield under high nitrogen condition but also more robust to low nitrogen stress. In contrast to grain yield, the morpho-physiological traits did not exhibit significant degrees of change in response to the degree of nitrogen limitation applied in this study, and of the traits which did exhibit significant effects – such as leaf nitrogen content and specific leaf area – the effects of treatment and genotype were largely independent of each other (Figure 2). While changes in chlorophyll concentration were quantifiable using both handheld chlorophyll concentration meter and hyperspectral reflectance data, plants in the nitrogen limited field were not visibly chlorotic (personal observation). Metabolite abundance was characterized for a subset of sorghum genotypes in both conditions in an attempt to identify other phenotypes with potential value to predict how the grain yield of different sorghum varieties will respond to nitrogen limitation. The overall pattern of metabolite abundances did not exhibit substantial differences between nitrogen limited and non-nitrogen limited conditions (Fig. 3a-c). This is consistent with both the limited degree of change observed for morpho-physiological traits and the goal of imposing a degree of nitrogen limitation sufficient to alter fitness/grain yield but not so severe that it dramatically altered plant growth.

While overall differences in metabolite abundance between conditions were modest, the metabolites that did exhibit significant differences between treatments were consistent with expectations for nitrogen limited grown plants. Decreases in the abundance of many amino acids were observed (Fig. 3d; Table S4). Consistent with reports from studies of nitrogen deficit experiments in seedlings and adult maize leaves (Amiour *et al*., 2012), sorghum roots (Sheflin *et al*., 2019), and maize, sorghum, and *Paspalum vaginatum* seedlings (Sun *et al*., 2021). Disturbance in serine metabolism was previously found to play key role in limiting maize yield under low nitrogen conditions (Cañas *et al*., 2012). In addition, serine plays an important role in photorespiration (Maurino and Peterhansel, 2010), although in plants utilizing the C4 photosynthetic pathway, including both maize and sorghum this pathway is much less active than in plants utilizing the C3 photosynthetic pathway. Unfortunately, we did not observe significant variation in the degree of decreased serine abundance observed among sorghum genotypes (Fig. 5c) suggesting that, while genetically controlled diversity for this trait may still be discovered in profiling of a larger panel of diverse sorghum lines under nitrogen limited and non nitrogen limited conditions, if no such diversity is found may prove impossible to reduce this response to nitrogen constrained growth via conventional breeding and selection strategies. In contrast, while the abundance of glutamic acid also declined in nitrogen limited conditions, the degree of decline varied significantly among sorghum genotypes (Fig. 5d). In previous field studies of maize, the abundance of glutamic acid was negatively correlated with yield under heat and water stress but not under control conditions (Obata *et al*., 2015). We observed a similar negative correlation between glutamic acid abundance and yield under both control and nitrogen limited conditions (Fig. 6). This result highlight potential importance of glutamic acid metabolism on yield in C4 crops under stress conditions. Genes involved in glutamic acid metabolism were enriched among those exhibiting differential mRNA expression between older maize inbreds (pre-1960s) and maize inbreds developed and selected by breeders in the modern era (Xu *et al*., 2022). These observations suggest that glutamic acid metabolism may already have been an indirect target of selection during crop improvement in maize. If so, the data presented here suggest that glutamic acid may also represent an interesting metabolic marker when selecting for better performing sorghum genotypes although further validation is certainly needed. Overall, our results highlight that grain yield in sorghum, unlike many morpho-physiological traits, exhibits substantial variability of genotype specific responses to long term low severity nitrogen deficit stress. Differences in the eight morpho-physiological traits scored in this study explained only ~ 9% of variance in yield. Metabolic responses to long term low severity nitrogen deficit stress exhibited a higher proportion of variability explained by genotype specific responses than did morpho-pysiological traits and a number of individual metabolites were associated with yield variation under one or both nitrogen treatments. It may be possible to build predictive models using metabolite abundance to estimate which sorghum genotypes will exhibit greater or lesser decreases in yield in response to nitrogen deficit, however data from a larger number of genotypes grown across multiple sites will be necessary to train and evaluate such models. Large scale metabolic profiling will likely require targeted metabolomics using feature selection approaches to identify an informative subset of the metabolites profiled in this study.

## Acknowledgements

This project was supported by U.S. Department of Energy, Grant no. DE-SC0020355 to JCS, the National Science Foundation under grant OIA-1826781, USDA-NIFA under the AI Institute: for Resilient Agriculture, Award No. 2021-67021-35329 and the Foundation for Food and Agriculture Research Award No. 602757. This project was enabled by the work of the the Proteomics & Metabolomics Facility (RRID:SCR_021314) of Nebraska Center for Biotechnology at the University of Nebraska-Lincoln, utilizing instrumentation are supported by the Nebraska Research Initiative.

## Author contributions

J. C. S., M. W. G., Y. G. and S.A. conceived and designed the study. M. W. G., M. Z., H. J., N. K. W., A. A., M. N. and S. A. collected data. M. W. G. analysed the data with guidance from J. C. S. and S. A. M. W. G and J. C. S. wrote initial draft of manuscript with input from S. A. All authors approved the final version of the manuscript.

## Data availability

Hyperspectral data are available from https://ecosis.org/package/eb492231-a217-4aaa-9857-dae642a85ed3. Raw metablomic data are available from Metabolight under accession number: MTBLS4951.

---

**Table S1. Plot-level values of morpho-physiological traits**. Provided as Excel file.

**Table S2. Raw ground truth and spectral data used to build PLSR models and predict three traits**. Provided as csv file.

**Table S3.** Summary of the PLSR results

| Trait | n<sub>LV</sub> | Training |  | Validation |  |
| --- | --- | --- | --- | --- | --- |
|  |  | R<sup>2</sup> | RMSE | R<sup>2</sup> | RMSE |
| CHL | 19 | 0.82 | 48.73 | 0.83 | 46.22 |
| SLA | 10 | 0.62 | 6.47 | 0.76 | 5.97 |
| N | 16 | 0.66 | 0.29 | 0.46 | 0.34 |
| P | 10 | 0.18 | 0.06 | 0.25 | 0.07 |
| K | 10 | 0.34 | 0.34 | 0.22 | 0.37 |

**Table S4. Summary statistics for 145 annotated metabolites**.

**Table S5. Correlation values between yield and 145 metabolites**.

**Figure S1.**
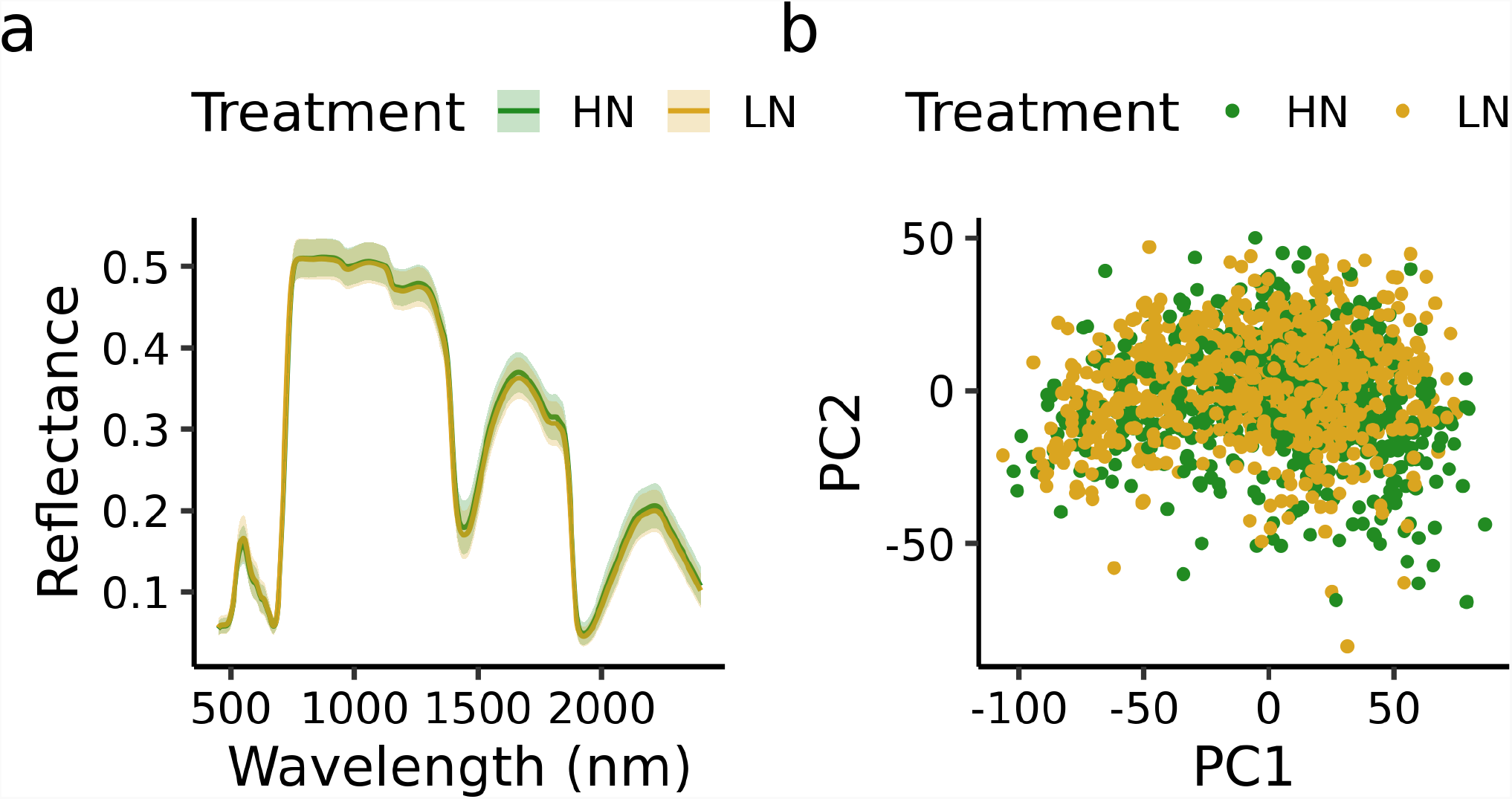
**a** The mean leaf hyperspectral of the sorghum plants from high nitrogen(HN; green) and low nitrogen(LN; yellow). The bounding envelopes are the standard deviation. **b** Principal component score for individual plat (PC1 vs. PC2).

**Figure S2.**
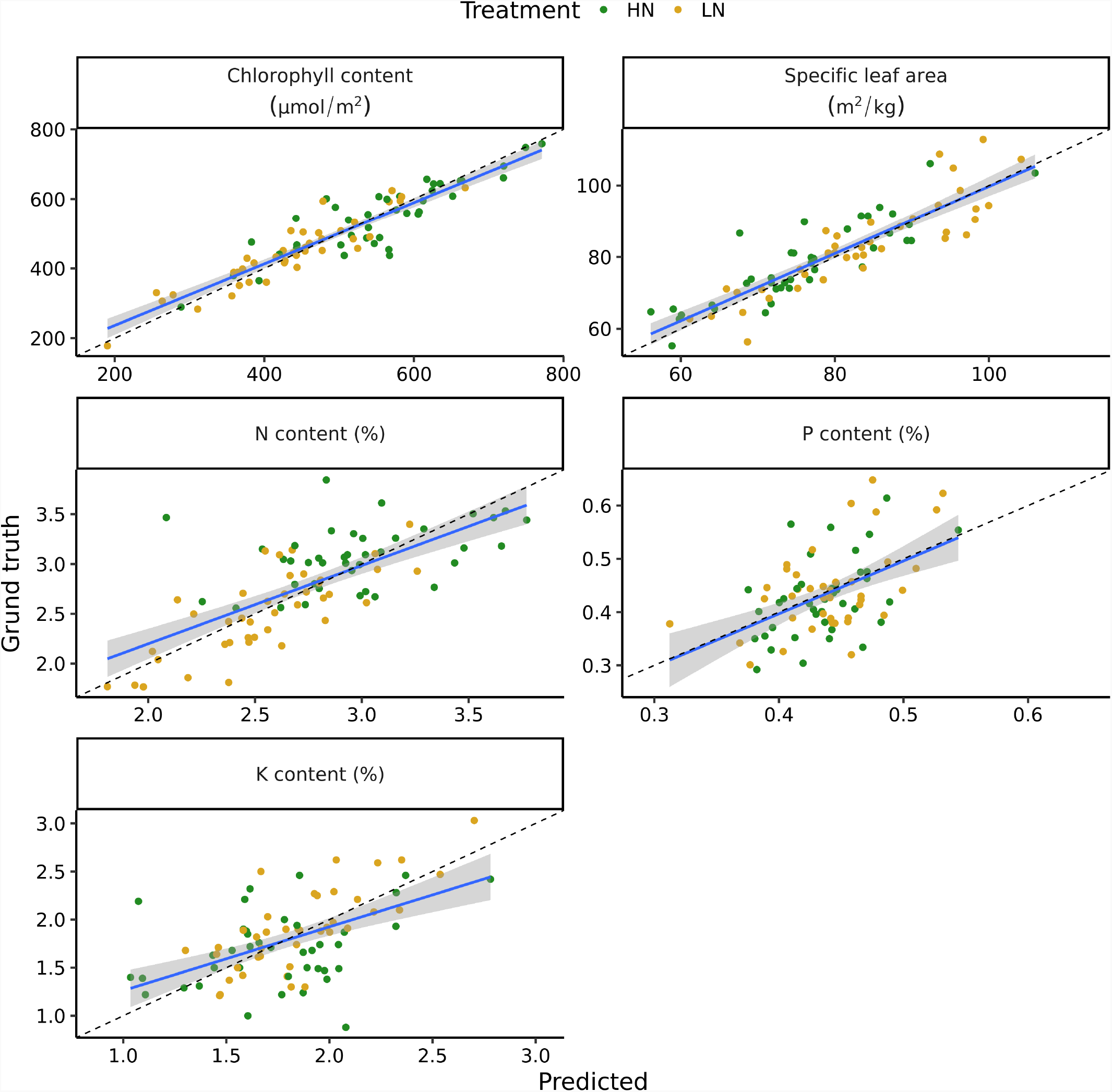
Scatter plots of ground truth and predicted values for training set sorghum leaves. Statistics for prediction can be found in Table S3. HN - high nitrogen, LN - low nitrogen.

**Figure S3.**
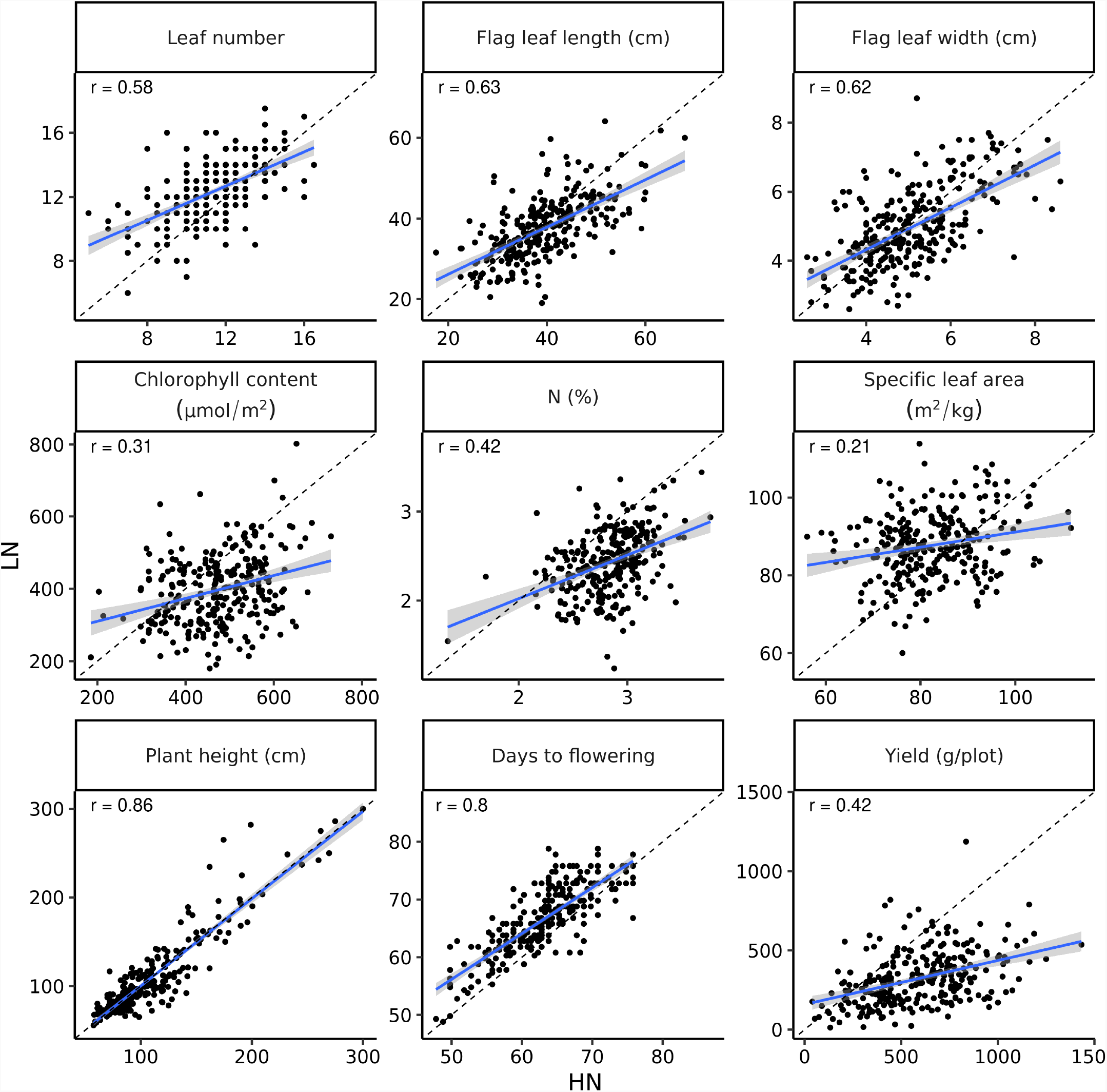
Scatter plot of genotype mean of morpho-physiological traits between two nitrogen conditions. r indicates Pearson correlation value. HN - high nitrogen, LN - low nitrogen.

**Figure S4.**
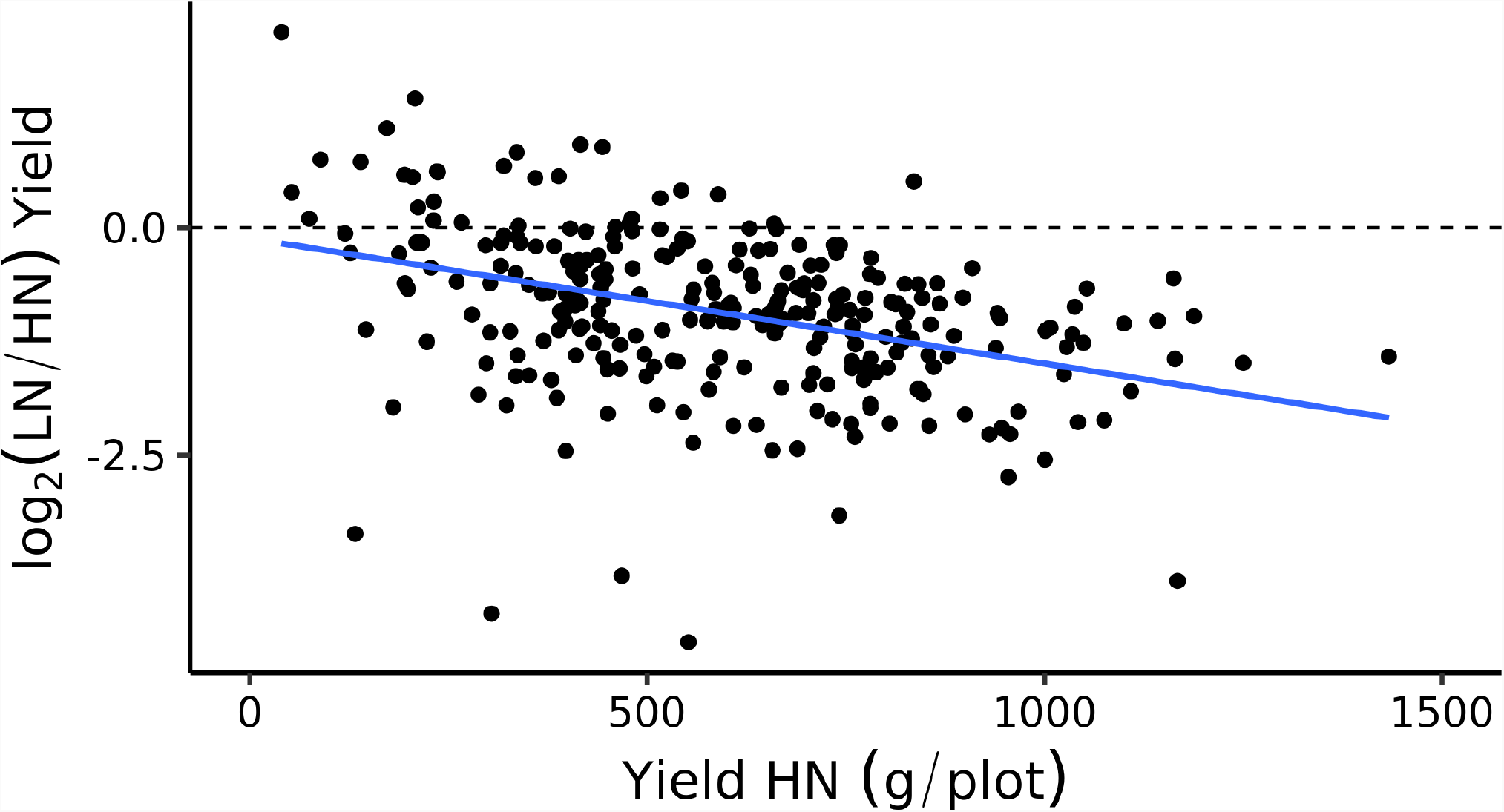
Plasticity in yield under low nitrogen (LN) stress versus yield at control (HN - high nitrogen) conditions. The dotted line indicates zero difference (no low nitrogen effect), while the solid blue line is the fitted regression. Each dot represents a genotype mean.

**Figure S5.**
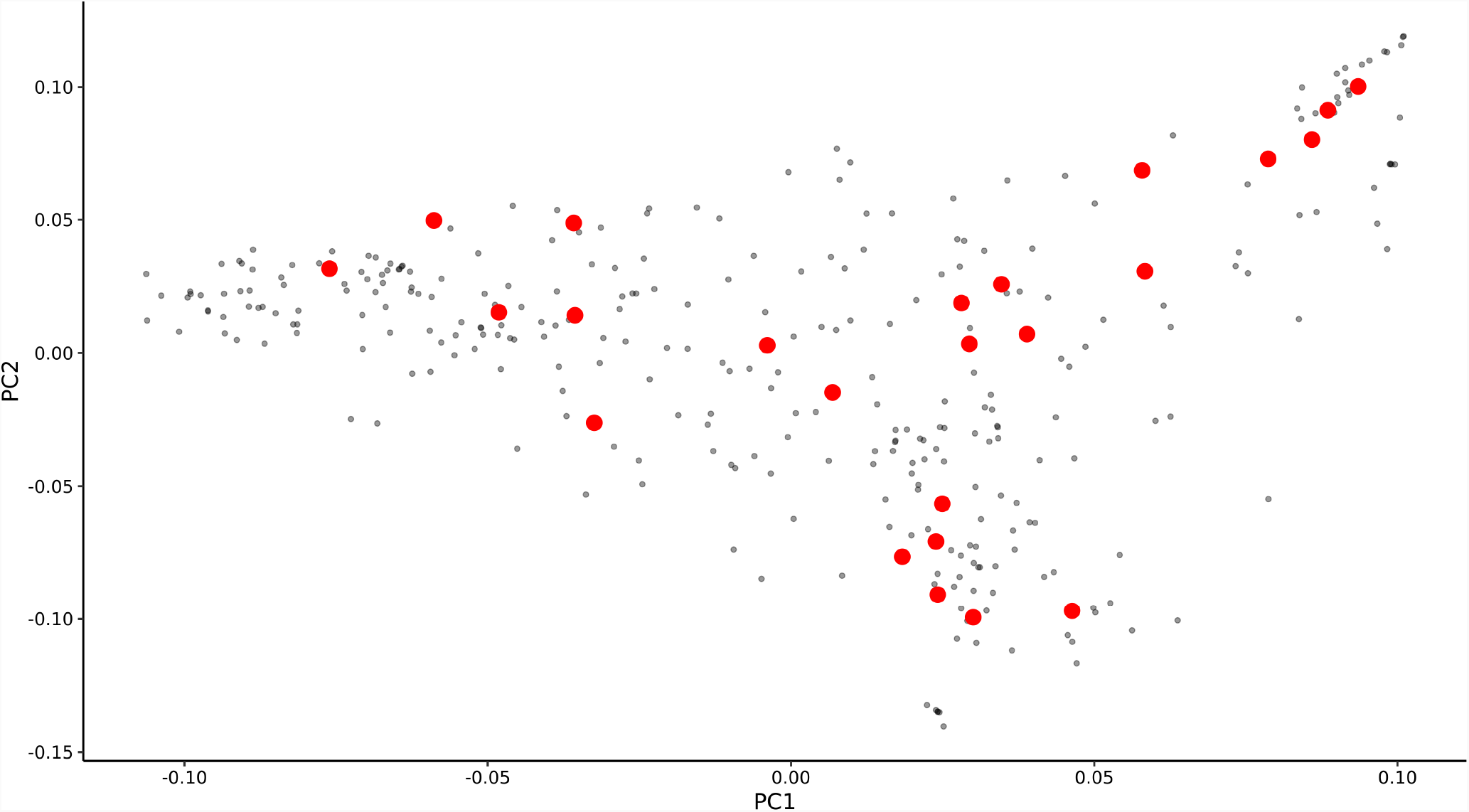
First two principle component from PCA based on SNPs from Miao *et al*. (2020). Each dot represent single genotype and red dots marked genotypes selected to metabolomic analysis.

**Figure S6.**
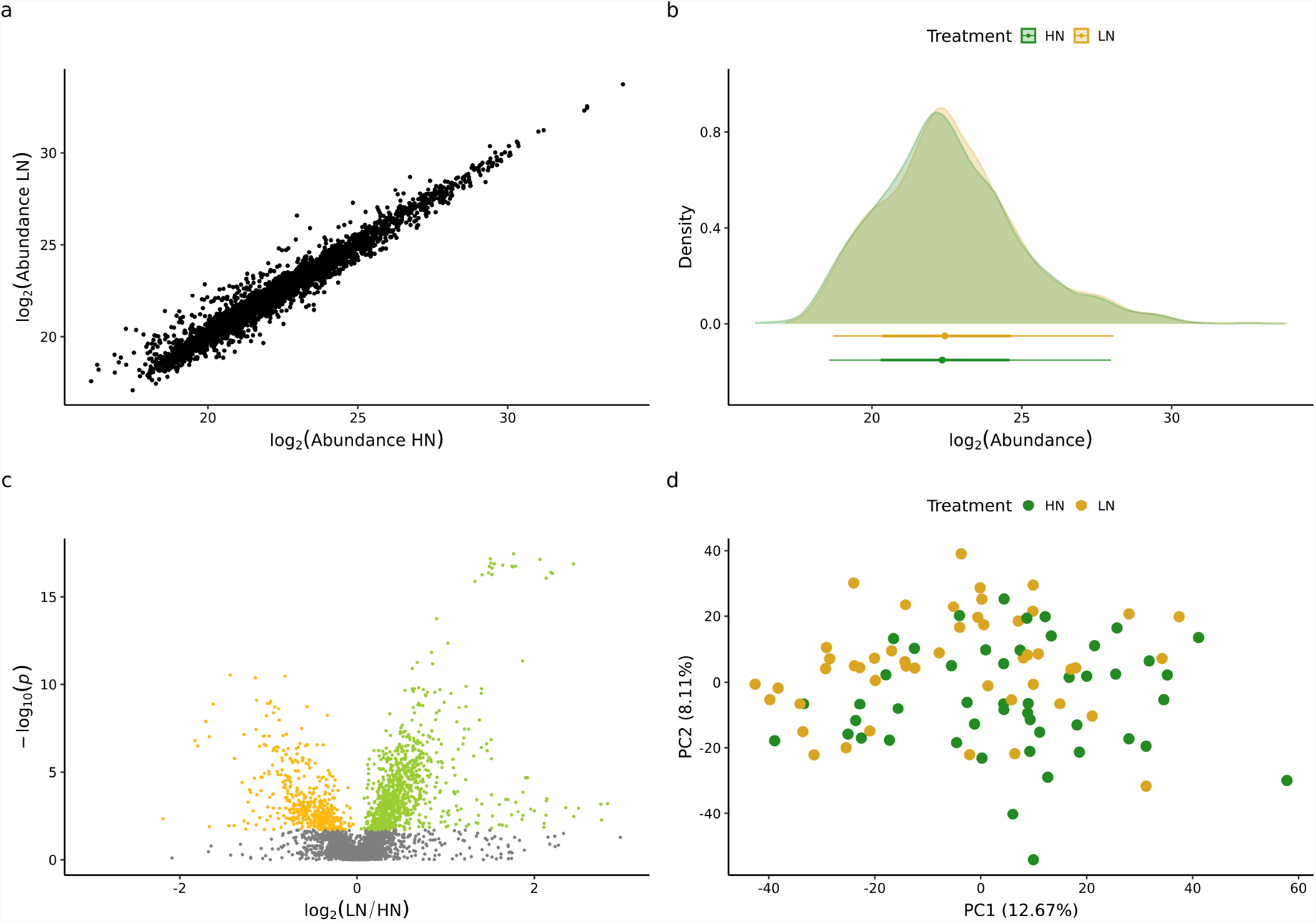
Metabolomics profiling in 24 sorghum genotypes across two nitrogen conditions based 3,496 identified compounds. **a** Scatter plot of abundance of 3,496 identified compounds across two treatment conditions. **b** Distribution of 3,496 identified compounds treatment conditions **c**. Volcano plot showing the downregulated (yellow) and upregulated (green) metabolites under low nitrogen (LN) conditions compare to high nitrogen (HN). **d** First two principle components (PC) from PCA based on 3,496 identified compounds. Values in bracket indicate amount of variance explained by each component. HN - high nitrogen, LN - low nitrogen.

**Figure S7.**
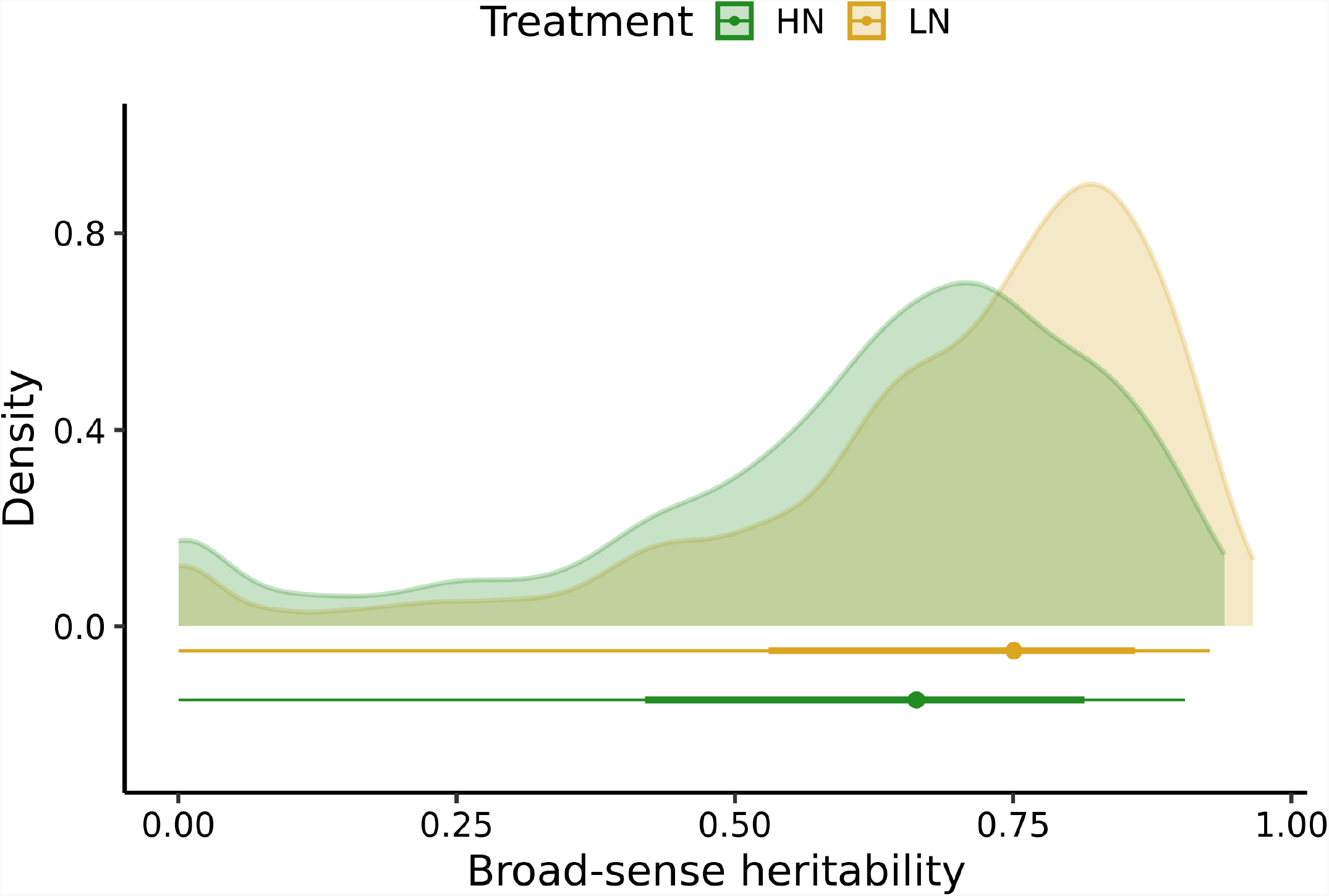
Distribution of broad-sense heritability (H^2^) values for 3,496 identified compounds in two treatment condition. HN - high nitrogen, LN - low nitrogen.

**Figure S8.**
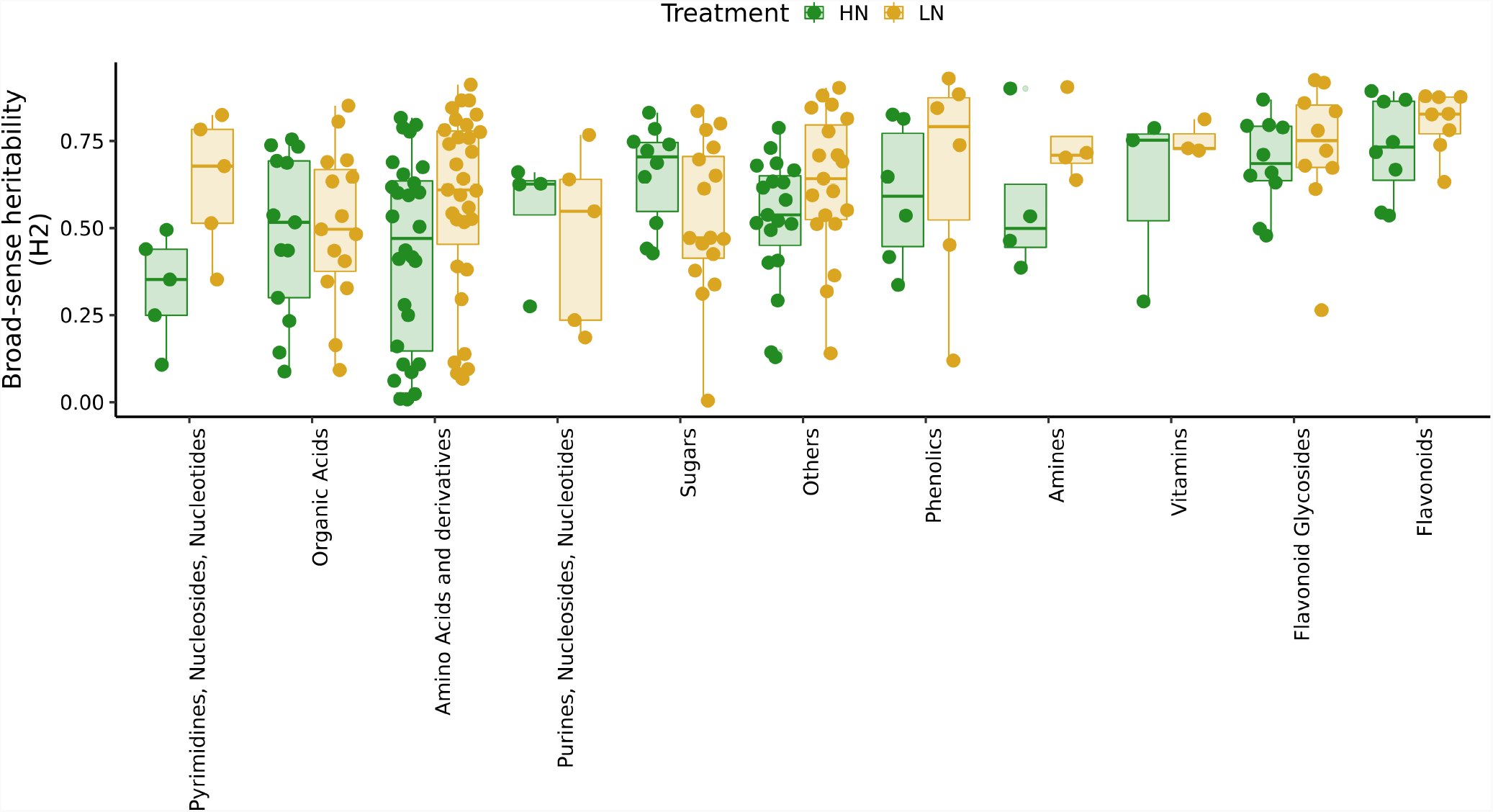
Broad-sense heritability (H^2^) values for 145 annotated metabolites across 11 classes. Each dot indicate a single metabolite. HN - high nitrogen, LN - low nitrogen.

**Figure S9.**
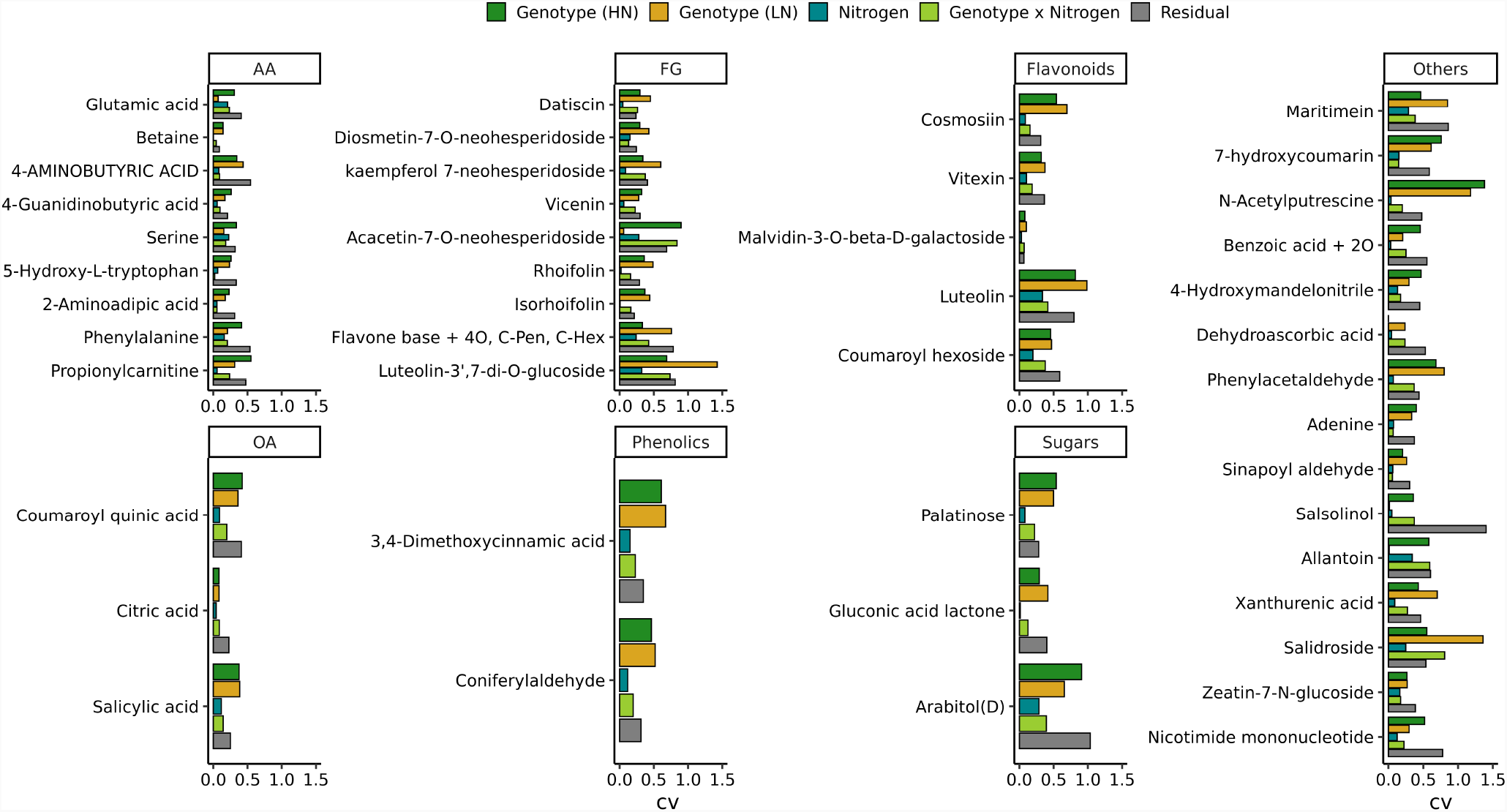
Coefficient of variation for 45 metabolites (CV; the estimated variance divided by the squared mean of the respective trait).

**Figure S10.**
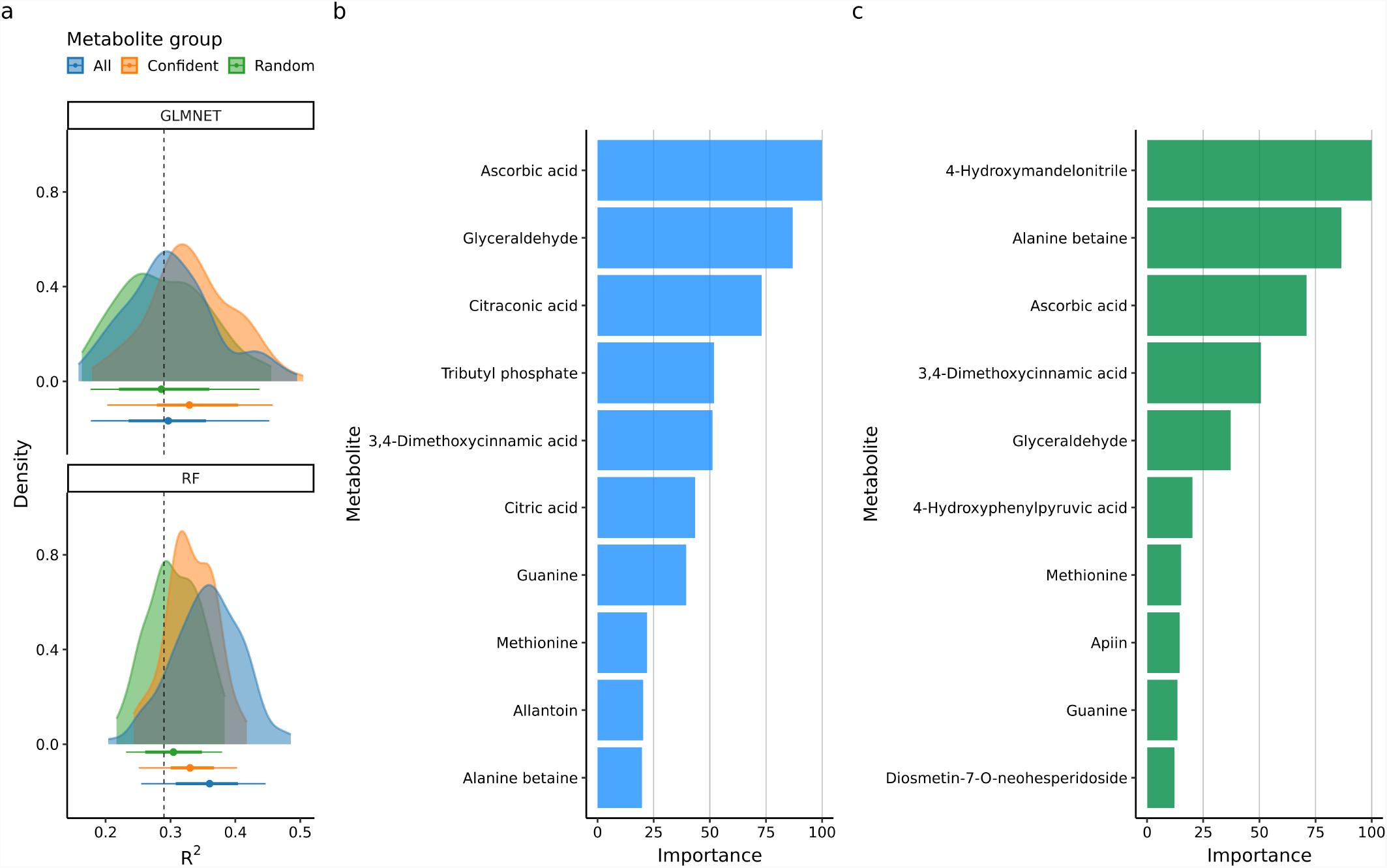
**a** Yield accuracy prediction based on three metabolites sets: all identified metabolites (n=3,496), metabolites with confident annotation (n=145) and the same number of metabolites with unknown annotation (n=145). R^2^ were obtained from 100x repeated five-fold cross-validation. Dashed lines indicated R^2^ values from regression based on treatment conditions. GLMNET - elastic-net regression, RF - random forest. Imporance values based on permutation for GLMNET(**b**) and RF(**c**).

## References

Amiour, N., S. Imbaud, G. Clément, N. Agier, M. Zivy, B. Valot, T. Balliau, P. Armengaud, I. Quilleré, R. Cañas, et al., 2012 The use of metabolomics integrated with transcriptomic and proteomic studies for identifying key steps involved in the control of nitrogen metabolism in crops such as maize. Journal of Experimental Botany 63: 5017–5033.

Banerjee, B. P., S. Joshi, E. Thoday-Kennedy, R. K. Pasam, J. Tibbits, M. Hayden, G. Spangenberg, and S. Kant, 2020 High-throughput phenotyping using digital and hyperspectral imaging-derived biomarkers for genotypic nitrogen response. Journal of Experimental Botany 71: 4604–4615.

Bates, D., M. Mächler, B. Bolker, and S. Walker, 2015 Fitting linear mixed-effects models using lme4. Journal of Statistical Software 67: 1–48.

Benjamini, Y. and Y. Hochberg, 1995 Controlling the false discovery rate: a practical and powerful approach to multiple testing. Journal of the Royal statistical society: series B (Methodological) 57: 289–300.

Brommer, J. E., 2013 Variation in plasticity of personality traits implies that the ranking of personality measures changes between environmental contexts: calculating the cross-environmental correlation. Behavioral Ecology and Sociobiology 67: 1709–1718.

Cañas, R. A., I. Quilleré, A. Gallais, and B. Hirel, 2012 Can genetic variability for nitrogen metabolism in the developing ear of maize be exploited to improve yield? New Phytologist 194: 440–452.

Casa, A. M., G. Pressoir, P. J. Brown, S. E. Mitchell, W. L. Rooney, M. R. Tuinstra, C. D. Franks, and S. Kresovich, 2008 Community resources and strategies for association mapping in sorghum. Crop Science 48: 30–40.

Cormier, F., S. Faure, P. Dubreuil, E. Heumez, K. Beauchêne, S. Lafarge, S. Praud, and J. Le Gouis, 2013 A multi-environmental study of recent breeding progress on nitrogen use efficiency in wheat (triticum aestivum l.). Theoretical and Applied genetics 126: 3035–3048.

de Jong, M., H. Tavares, R. K. Pasam, R. Butler, S. Ward, G. George, C. W. Melnyk, R. Challis, P. X. Kover, and O. Leyser, 2019 Natural variation in arabidopsis shoot branching plasticity in response to nitrate supply affects fitness. PLoS genetics 15: e1008366.

Erisman, J. W., M. A. Sutton, J. Galloway, Z. Klimont, and W. Winiwarter, 2008 How a century of ammonia synthesis changed the world. Nature geoscience 1: 636–639.

Foley, J. A., N. Ramankutty, K. A. Brauman, E. S. Cassidy, J. S. Gerber, M. Johnston, N. D. Mueller, C. O’Connell, D. K. Ray, P. C. West, et al., 2011 Solutions for a cultivated planet. Nature 478: 337–342.

Friedman, J., T. Hastie, and R. Tibshirani, 2010 Regularization paths for generalized linear models via coordinate descent. Journal of Statistical Software 33: 1–22.

Gao, K., F. Chen, L. Yuan, F. Zhang, and G. Mi, 2015 A comprehensive analysis of root morphological changes and nitrogen allocation in maize in response to low nitrogen stress. Plant, cell & environment 38: 740–750.

Ge, Y., A. Atefi, H. Zhang, C. Miao, R. K. Ramamurthy, B. Sigmon, J. Yang, and J. C. Schnable, 2019 High-throughput analysis of leaf physiological and chemical traits with vis–nir–swir spectroscopy: a case study with a maize diversity panel. Plant methods 15: 1–12.

Hakeem, K. R., A. Ahmad, M. Iqbal, S. Gucel, and M. Ozturk, 2011 Nitrogen-efficient rice cultivars can reduce nitrate pollution. Environmental Science and Pollution Research 18: 1184–1193.

Kant, S., Y.-M. Bi, and S. J. Rothstein, 2011 Understanding plant response to nitrogen limitation for the improvement of crop nitrogen use efficiency. Journal of experimental Botany 62: 1499–1509.

Kuhn, M., 2008 Building predictive models in r using the caret package. Journal of statistical software 28: 1–26.

Lê, S., J. Josse, and F. Husson, 2008 FactoMineR: A package for multivariate analysis. Journal of Statistical Software 25: 1–18.

Liland, K. H., B.-H. Mevik, and R. Wehrens, 2021 pls: Partial Least Squares and Principal Component Regression. R package version 2.8-0.

Liu, Y., H. Wang, Z. Jiang, W. Wang, R. Xu, Q. Wang, Z. Zhang, A. Li, Y. Liang, S. Ou, et al., 2021 Genomic basis of geographical adaptation to soil nitrogen in rice. Nature 590: 600–605.

Malthus, T. R., 1798 An essay on the principle of population, as it affects the future imporvement of society, with remarks on the speculations of Mr. Godwin, M. Condorcet, and other writers. J. Johnson, London.

Maurino, V. G. and C. Peterhansel, 2010 Photorespiration: current status and approaches for metabolic engineering. Current opinion in plant biology 13: 248–255.

Miao, C., Y. Xu, S. Liu, P. S. Schnable, and J. C. Schnable, 2020 Increased power and accuracy of causal locus identification in time series genome-wide association in sorghum. Plant physiology 183: 1898–1909.

Obata, T., S. Witt, J. Lisec, N. Palacios-Rojas, I. Florez-Sarasa, S. Yousfi, J. L. Araus, J. E. Cairns, and A. R. Fernie, 2015 Metabolite profiles of maize leaves in drought, heat, and combined stress field trials reveal the relationship between metabolism and grain yield. Plant Physiology 169: 2665–2683.

R Core Team, 2021 R: A Language and Environment for Statistical Computing. R Foundation for Statistical Computing, Vienna, Austria.

Ramankutty, N., Z. Mehrabi, K. Waha, L. Jarvis, C. Kremen, M. Herrero, and L. H. Rieseberg, 2018 Trends in global agricultural land use: implications for environmental health and food security. Annual review of plant biology 69: 789–815.

Raun, W. R. and G. V. Johnson, 1999 Improving nitrogen use efficiency for cereal production. Agronomy journal 91: 357–363.

Rothstein, S. J., 2007 Returning to our roots: making plant biology research relevant to future challenges in agriculture. The Plant Cell 19: 2695–2699.

Sheflin, A. M., D. Chiniquy, C. Yuan, E. Goren, I. Kumar, M. Braud, T. Brutnell, A. L. Eveland, S. Tringe, P. Liu, et al., 2019 Metabolomics of sorghum roots during nitrogen stress reveals compromised metabolic capacity for salicylic acid biosynthesis. Plant direct 3: e00122.

Sumner, L. W., A. Amberg, D. Barrett, M. H. Beale, R. Beger, C. A. Daykin, T. W.-M. Fan, O. Fiehn, R. Goodacre, J. L. Griffin, et al., 2007 Proposed minimum reporting standards for chemical analysis. Metabolomics 3: 211–221.

Sun, G., N. Wase, S. Shu, J. Jenkins, B. Zhou, C. Chen, L. Sandor, C. Plott, Y. Yoshinga, C. Daum, et al., 2021 The genome of stress tolerant crop wild relative paspalum vaginatum leads to increased biomass productivity in the crop zea mays. bioRxiv.

Thurber, C. S., J. M. Ma, R. H. Higgins, and P. J. Brown, 2013 Retrospective genomic analysis of sorghum adaptation to temperate-zone grain production. Genome biology 14: 1–13.

Tsugawa, H., T. Cajka, T. Kind, Y. Ma, B. Higgins, K. Ikeda, M. Kanazawa, J. VanderGheynst, O. Fiehn, and M. Arita, 2015 Ms-dial: data-independent ms/ms deconvolution for comprehensive metabolome analysis. Nature methods 12: 523–526.

Wickham, H., M. Averick, J. Bryan, W. Chang, L. D. McGowan, R. François, G. Grolemund, A. Hayes, L. Henry, J. Hester, et al., 2019 Welcome to the tidyverse. Journal of Open Source Software 4: 1686.

Wright, M. N. and A. Ziegler, 2017 ranger: A fast implementation of random forests for high dimensional data in C++ and R. Journal of Statistical Software 77: 1–17.

Xu, G., J. Lyu, T. Obata, S. Liu, Y. Ge, J. C. Schnable, and J. Yang, 2022 A historically balanced locus under recent directional selection in responding to changed nitrogen conditions during modern maize breeding. bioRxiv.

Zhang, X., E. A. Davidson, D. L. Mauzerall, T. D. Searchinger, P. Dumas, and Y. Shen, 2015 Managing nitrogen for sustainable development. Nature 528: 51–59.

